# APP^NL-F^ knock-in Alzheimer’s Disease model mice reveal etiological convergence of aging and amyloidosis

**DOI:** 10.64898/2026.01.28.702259

**Authors:** Matthieu Prieur, Anne-Laure Hemonnot-Girard, Valentin Beck, Valentin Garcia, Thanawat Phuadraksa, Nathalie Linck, Manon Leportier, Anaïs Vignon, Dany Severac, Véronique Perrier, François Rassendren, Marc Dhenain, Takashi Saito, Takaomi Saido, Hélène Hirbec

**Author notes:** These two authors contributed equally to this work (Matthieu Prieur, Anne-Laure Hemonnot-Girard). Correspondence: Hirbec Hélène.

## Abstract

Alzheimer’s disease (AD) is the leading cause of dementia worldwide, with a steadily increasing prevalence. Despite the advent of some plaque-degrading therapies, early intervention strategies are still suboptimal, largely due to an incomplete understanding of how early pathological events of amyloid-β (Aβ) plaque deposition entail or accompany the neuroinflammatory processes that ensue. The APP^NL-F^ mouse line is a second-generation endogenous-promoter knock-in AD model that represents a valuable tool to dissect disease mechanisms and evaluate therapeutic strategies. However, the early prodromal stage remains poorly characterized. Here we performed in-depth analyses in both homozygous (APP^NL-F/NL-F^) and heterozygous (APP^NL-F/WT^) mice at 3, 6, 9 and 12 months of age. Among a battery of behavioural tests, the first phenotype was observed at 9 months with defects in spatial memory. Using hypersensitive MSD-ELISA technology assays we quantified distinct Aβ species and found that Aβ42 oligomers, protofibrils, and fibrillar aggregates were detectable as early as 6 months of age in homozygous APP^NL-F/NL-F^ mice. By 9 months, their Aβ42 levels increased markedly and overt Aβ plaques were detected, histologically associated with recruited glial cells. Targeted RT-qPCR analysis of neuroinflammation-related genes in the cortex also identified 9 months as a molecular tipping point in these ‘middle aged’ APP^NL-F^ mice. To characterize the etiological signal transduction at the cellular level, we isolated microglia, the brain resident immune cells, whose contribution to AD pathogenesis is now well established. Sequencing around 5,000 individual cells from both 12-month-old APP^NL-F/NL-F^ and APP^WT/WT^ CD11b+ myeloid cells revealed that the buildup of amyloidosis was associated with an accelerated shift of microglia from a homeostatic toward a senescent-like state. Together, these findings highlight the 6-12 month period of the APP^NL-F/NL-F^ model as a powerful system to study the interdependence between microglial senescence and amyloidosis in driving AD progression.

## Introduction

Alzheimer’s disease (AD) is the most prevalent form of dementia and represents a major and unresolved medical challenge. Despite decades of intensive research and the recent development of passive immunisation strategies ^1^, truly effective and well-tolerated disease-modifying treatments remain elusive. This therapeutic gap reflects persistent uncertainties in our understanding of the molecular mechanisms that initiate and propel the disease, particularly during its early phases. Neuropathologically, AD is characterized by extracellular accumulation of amyloid-β (Aβ) peptides, intracellular aggregation of hyperphosphorylated Tau, and a neuroinflammatory response ^2–4^. Among these, Aβ deposition is widely regarded as an early event that sets the stage for Tau pathology, which in turn leads to neurodegeneration ^5^. Advancing age represents the main risk factor for AD, yet the molecular and cellular processes linking aging and Aβ deposition remain incompletely understood.

In AD, mounting evidence points to the involvement of glial cells, particularly microglia and astrocytes, which enter into reactive states and contribute to both pathology and neuronal dysfunction ^6^. Among these reactive microglial states, Disease Associated Microglia (DAM) are a distinct transcriptional and functional cell type, first identified in AD but that also emerge in response to various neurodegenerative and chronic diseases of the central nervous system ^7,8^. During aging, microglia progressively acquire features of senescence, characterized by transcriptional changes, reduced surveillance capacity, impaired phagocytosis and excessive inflammatory responses ^9^. However, the extent to which the senescence trajectories of glial cells affect or depend on rising amyloid burden to trigger maladaptive states remains poorly defined, representing a key mechanistic gap in our understanding of AD pathogenesis. Clarifying any potential interdependence between amyloidosis and microglial senescence is critical to determining if the latter has significant therapeutic potential.

Probing the interdependence of amyloidosis and aging requires experimental models in which amyloid accumulation and cellular senescence progress synchronically. First-generation AD models, based on transgenic overexpression of human mutant APP and presenilin-1 (PS1) ^10^, have been instrumental in advancing the understanding of Aβ impact on the brain. However random transgene integration suffers from potential off-target or artifact effects and non-physiological overexpression levels potential off-target or artifact effects ^11^. In particular, most studies were conducted on relatively young animals, that already displayed high amyloid load. This diminishes their translational relevance as they fail to reveal the impact of amyloid in an aging brain and organism. More refined second-generation knock-in mouse models have thus been developed to overcome these issues and more faithfully reproduce key aspects of human AD by precisely inserting Aβ-producing APP mutants under the control of the mouse APP promoter ^11^. Among them, the APP^NL-F^ knock-in mouse is increasingly considered as one of the most pathophysiologically relevant models of amyloidosis. By introducing the Swedish (NL) and Iberian (F) mutations into the endogenous humanized *App* locus, this line promotes the amyloidogenic processing of APP while preserving a wild-type human Aβ sequence, resulting in a gradual accumulation of amyloid pathology ^12^. Importantly, the midlife onset of Aβ deposition in APP^NL-F^ mice uniquely positions this model to address an unresolved question in the field: how aging-related processes interact with the earliest stages of amyloid-driven AD pathogenesis.

In this study, we systematically investigated early-stage pathology in the APP^NL-F^ mouse model, spanning 3 to 12 months of age, in both heterozygous and homozygous animals. Our analysis integrated multiple dimensions of pathology, including behavioural alterations, molecular changes in Aβ species, and an in-depth assessment of neuroinflammation, with a particular focus on the temporal emergence of microglial vulnerability. We show that most pathological alterations emerge at around 9 months in our experimental conditions. Through single-cell RNA sequencing of 12-month-old mutant and wild-type brains, we uncovered profound transcriptomic remodelling and identified senescent-like microglia as highly sensitive to amyloid pathology. Our findings thus highlight a mechanistic convergence between amyloid accumulation and aging-related microglial dysfunction, establishing the APP^NL-F^ model as a powerful platform for investigation and therapeutic targeting of the early intersection of amyloidosis and aging.

## Materials and Methods

### Animals

Human amyloid precursor protein (hAPP) knock-in (KI) mice carrying the Swedish (*KM670/671NL)* and Iberian (I716F) mutations (App^NL-F^) were originally generated by Dr Saito on a C57BL/6J background ^12^. In this study, we analyzed three genotypes: APP^WT/WT^ (WT), APP^NL-F/WT^ (NL-F/WT), and APP^NL-F/NL-F^ (NL-F/NL-F), where WT denotes the endogenous murine APP allele. Genotyping of the App^NL-F^ allele was performed by quantitative PCR (qPCR). Throughout the manuscript, APP^NL-F^ designates the KI line, irrespective of genotype. All mice were maintained on a C57BL6/J background and generated through breeding of APP^NL-F/WT^ mice in the “Specific Pathogen-Free” animal facility of the Institute of Functional Genomics (IGF, Montpellier, France; Ministry of Agriculture authorization N° D34-172-13). Mice were housed at 22°C, in a humidity of 55% (± 10%), under a 12-hour light/dark cycle with ad libitum access to food and water.

All experiments were conducted in compliance with European Union guidelines (Directive 2010/63/EU) and institutional guidelines for the care and use of laboratory animals. The animal experimentation protocols used in this study were approved by the Ethics Committee for Animal Experimentation of Languedoc Roussillon (CEEA-LR; APAFiS#5252).

Experiments were performed on animals aged 3, 6, 9, and 12 months, both males and female were tested.

### Behavioral Tests

Cognitive performance was assessed in a series of non-invasive tests. To minimize stress, animals underwent a 5-day habituation protocol with the experimenter one week prior to testing. Behavioral assays were performed in the following order: Openfield, then Barnes Maze, progressing from the least to the most cognitively demanding task. Between each trial, apparatuses were cleaned with 20% ethanol followed by water to eliminate olfactory cues. Exploration-based tests (Openfield and Y-maze) were performed under 50 lux non-aversive light. Animal movements were recorded and analysed using a video tracking system (Ethotrack software, Innovation Net).

The ***OpenField*** setup consisted of a white plastic arena measuring 44 x 44 x 30 cm. At the start of each trial, the mouse was placed in the center of the arena and allowed to explore freely for at least 10 minutes. The central zone was defined as the inner 50% of the total arena area. The total distance traveled and the distance traveled within the central zone were automatically quantified. Each mouse was tested only once.

The ***Barnes Maze*** apparatus consisted of a circular platform (92 cm in diameter, elevated 105 cm above the ground) with 20 equally spaced peripheral holes (5 cm in diameter). A dark “home box” was placed beneath one designated “target” hole, providing both refuge and reward (1 min in an enclosed, dark space). Visual cues were positioned on the walls of the testing room to support spatial navigation. The maze was illuminated at 350-370 lux to create a mildly aversive environment, motivating the mouse to locate the “home box”.

The testing protocol began with a one-day habituation session, during which each mouse was allowed to explore the maze once for 60 seconds before being gently guided to the “home box” for 120 s rest period. This was followed by a 4-day <u>acquisition phase</u> (i.e. learning phase), during which mice learned to locate the “home box” using external visual cues. Each daily training session consisted of four 3-min trials separated by 15-min inter-trials intervals. During each trial, the mouse was allowed to freely explore the maze until finding the “home box”. Learning performance was assessed by measuring primary errors (i.e. number incorrect hole visits prior to locating the target) and primary latency (i.e. time to reach the target hole). <u>Spatial memory retention</u> was assessed 72 hours after the final learning session. During this probe trial, the “home box” was removed, and each mouse was allowed to explore the maze for 90 sec. The number of errors (visits to non-target holes) and the latency to reach the former target hole location were recorded. Mice were tested once only.

### Histological and Immunohistochemical Staining

#### Tissue Preparation

Following deep anesthesia, induced by intraperitoneal injection of Euthasol (2 µg/g, vetoEUT003AA, TVM), mice were transcardially perfused with phosphate-buffered saline (DPBS, D8537, Sigma-Aldrich). Brains were rapidly extracted and the hemispheres processed for distinct downstream applications. For histological analyses, one hemisphere was fixed in 4% paraformaldehyde (PFA, 15714, Electron Microscopy Sciences) for 2 h at room temperature (RT), followed by 24 h at 4°C. For transcriptomic or biochemical studies, the cortex of the second hemisphere was dissected and immediately stored at -80°C until further processing.

#### Histology and Immunohistochemistry

Coronal brain sections (30 µm thick) were obtained using a vibratome (VT1000S, Leica), with tissues blocks immersed in cold PBS during slicing. Sections were collected from regions of interest.

Dense amyloid plaques were visualized using Thiazine Red (ThzRed), a naphthol-based azo dye analog that binds to β-sheet structures (i.e., Like Thioflavin-S, it stains dense-core plaques but with a peak emission at 580 nm). Sections were incubated for 5 min in a 16 mg/L ThzRed solution (S570435, Sigma), then rinsed in PBS.

For immunohistochemistry, free-floating sections were permeabilized and blocked for 1 hour with PBS containing 10% donkey serum (Sigma-Aldrich, S30-100mL) and 0.3% Triton-X100 (Sigma, X100-100mL). Sections were then incubated at 4°C for 72 hours in the same blocking solution containing primary antibodies: anti-IBA1 to detect microglia (Abcam, AB178846, 1/2000), 6E10 to detect amyloid deposits (BioLegend, 803001, 1/500), or GFAP to detect astrocytes (Sigma, G3893, 1/1000). After thorough PBS washes, sections were incubated for 2 hours in blocking solution with appropriate secondary antibodies diluted in the blocking solution. Cell nuclei were stained with DAPI (B2883, Sigma). Sections were mounted onto glass slides using Fluorescence Mounting Medium using Dako (S3023, Dako).

#### Image acquisition and quantification

Images were taken with an AxioScan automated slide scanner (AxioScan, ZEISS) equipped with a 20x objective (magnification x20, numeric aperture 0.8, ZEISS). For each mouse, three rostral brain sections (i.e. at the level of the frontal cortex) and three caudal sections (i.e. at level of the hippocampus) were imaged entirely. The percentage of stained area in the region of interest was determined using Fiji software with specific macros: An average threshold was calculated for each mark, above which the analysis considered the mark to be positive. Regions of interest were manually drawn on each slice in the two brain regions studied (cerebral cortex and hippocampus).

### Transcriptomic Analyses

#### RNA Extraction

On the day of extraction, frozen tissues were homogenized at 4 m/s for 20 s using a FastPrep-24 Classic homogenizer (MPBio). Total RNA was isolated using the RNeasy Plus Mini Kit (Qiagen, 74136) following the instructions of the manufacturer. RNA concentration was measured in a NanoDrop2000 spectrophotometer (Thermo Fisher Scientific). RNA integrity was assessed with an Agilent 2100 Bioanalyzer (Agilent Technologies). All RNA used in this study had RNA integrity numbers (RIN) greater than 8.0.

#### qPCR

Total RNA was reverse-transcribed into cDNA using the iScript reverse transcription kit (Bio-Rad, 1708891), following the manufacturer’s protocol. qPCR was performed using gene-specific primers and SYBR Green for fluorescence-based amplification detection. Reactions were run on a LightCycler system (Roche), and Cq values were determined automatically using the second derivative method in the LightCycler software. Expression levels of the gene of interest were normalized to the housekeeping gene Hprt1, which was validated as a stable reference in the experimental conditions. Results were expressed as –ΔCq, calculated as: -ΔCq= -(Cq_GOI_ – Cq_Hprt1_). Primers used in the study were:

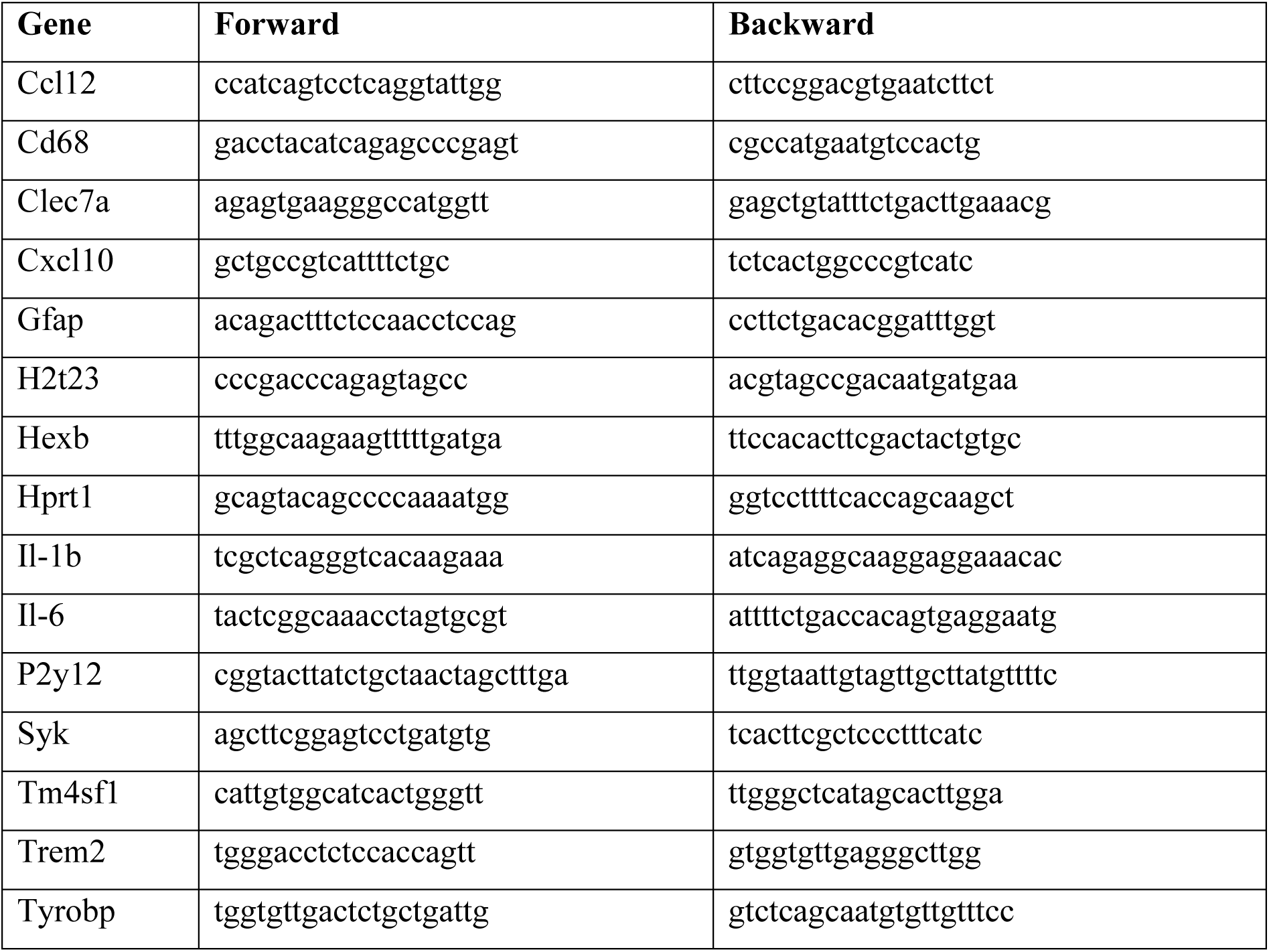

### Biochemical Analyses

#### Quantification of Aβ species

To minimize amyloid loss, all ELISA steps were carried out in LoBind Protein tubes (Eppendorf, 022431081). Cortical samples were weighed and homogenized at the ratio of 2.5 µl of buffer 1 per mg of tissue, with buffer 1 consisting of 50 mM Tris-HCl (Sigma, T2788) buffer containing, 1X Protease Inhibitor (PI) (ThermoFisher, A32961), 150 mM NaCl (Sigma, S5886) and 1% Triton X-100 (Sigma, 066K0089), pH 7.4. Homogenization was performed by sonication (7 pulses at 40% amplitude) using a Fischer Scientific FB120 sonicator (120 W, 20kHz, 230V); the samples were then centrifuged at 17,500 g, 4°C for 20 min. The resulting supernatant, corresponding to the Triton-soluble fraction (containing non-aggregated amyloids), was immediately stored at -80°C. The pellet was resuspended in 100 µl of 50 mM Tris-HCl containing 1X PI, 5M ganidine (Sigma, G7294), pH 7.4 and incubated at 800 rpm, 22°C for 3 h (Starlab, Mixer-HC). After centrifugation at 17,500g for 20 min at 4°C, the supernatant, representing the guanidine-Soluble fraction (containing aggregated amyloids), was collected and immediately stored at -80°C. Protein concentrations of both Triton-soluble and guanidine-soluble fractions were determined using the BCA assay. Samples were then diluted with Diluent 35 (MSD) to remain within the dynamic range of the assay. Levels of Aβ-40, Aβ-42 and Aβ-38 concentration were quantified using the MSD multiplex Aβ 6E10 kit (MSD, K15200E) following the manufacturer’s instructions. APP^WT/WT^ mice were used as controls, with all corresponding samples remaining below the detection limit. **Immunoblotting**: Cortical brain tissues were weighed and resuspended (10% w/v) in lysis buffer (140 mM KCl, 10 mM NaH_2_PO_4_, 1.7 mM KH_2_PO_4_, 1 mM EDTA, H20 milliQ) supplemented with 1X phosphatase inhibitor tablet (PhosSTOP, 04906845001, Roche) and 1X protease Inhibitor Cocktail (Mini cOmpleteTM,11836153001, Roche). Homogenization was performed in microbead-containing tubes using a ribolyzer apparatus (BioRad, Marnes La Coquette, France). Brain homogenates were supplemented with 2% SDS and centrifuged at 14,000 rpm for 20 min at 4°C to remove membrane debris. Protein concentrations in the supernatants were measured using a BCA protein assay (Pierce). All samples were normalized to the same protein concentration and diluted (1:1) in 2x loading buffer (4% SDS, 10% β-mercaptoethanol, 20% glycerol, 100 mM Tris-HCl pH 6.8, bromophenol blue). Samples were heated for 10 min at 90°C before loading on a 10% or 4–15% gradient gels (CriterionTM TGXTM Precast gels, Biorad). After electrophoresis and protein transfer, PVDF membranes were blocked in 5% nonfat milk-TBS-Tween solution and incubated ON at 4°C with one of the following primary antibodies: anti-BACE1 (Sigma Aldrich, SAB2100200); anti-IDE (Abcam, Ab32216); anti P-tau AT8 (Ser202, Thr205) (Life Technologies, MN1020); anti-total Tau AT5 (Life Technologies, AHB0042); anti-amyloid Aβ 6E10 (Biolegend, 803015). Protein loading was controlled using mouse anti-β-actin (Sigma Aldrich, A5441). After washing in TBS-T, the membranes were incubated for 1h at room temperature with either anti-mouse-HRP (Jackson Immuno Research, 115-035-003) or anti-rabbit-HRP (Sigma Aldrich, A6154) secondary antibody. Detection was performed using Luminata Crescendo substrate, and signals were acquired on a ChemiDoc™ MP Imager (Biorad) in signal accumulation mode. Quantification was carried out using image Lab software (version 6.0.1, Biorad) and normalized to β-actin signal.

#### Statistical Analyses of behavioural, immunohistochemistry and qPCR assays

Statistical analyses were performed using GraphPad Prism 9.0. Depending on the experimental design, one-way ANOVA, two-way ANOVA, Linear-mixed (LMM) models or Mann-Whitney tests were applied. When applicable, multiple comparisons were corrected using the False Discovery Rate (FDR) method. The specific statistical tests used are detailed in the main text and figure legends. Multivariate analyses, including Principal Component Analysis (PCA), and heatmap visualizations were conducted using dedicated R packages. Missing data accounted for less than 10% of the dataset and were imputed using the missMDA package in R. A p-value of <0.05 was considered statistically significant.

### Single cell experiments

#### Myeloid cell preparation and scRNA-seq analysis

Myeloid cell isolation was performed from 12-mo males APP^NL-F/NL-F^ (1 pool of 3 mice) and aged age-matched controls (APP^WT/WT^, 1 pool of 2 mice). In brief, PBS perfused brains were dissected on ice, cortices were isolated and cells were dissociated with the adult brain dissociation kit (ABDK Miltenyi Biotec, Germany) and the gentleMACS™ Octo Dissociator with Heaters (Miltenyi Biotec, Germany). The manufacturer’s protocol was followed with modifications. Specifically, 5% BSA was added to the dissociation mixture to enhance antigen preservation, and 10 µM Actinomycin D was included to minimize transcriptional activation during the process. After myelin removal, CD11B+ cells were isolated with magnetic bead-coupled anti-CD11B antibodies according to the manufacturer’s protocol (Miltenyi Biotec, Germany). Cell viability and presence of debris were evaluated using the Caysy cell counter (OLS).

Cell suspensions were loaded on a Chromium controller (10x Genomics, Pleasanton, CA, USA) to generate single-cell Gel Beads-in-Emulsion (GEMs). Single-cell RNA-seq libraries were prepared using Chromium Single cell 3’RNA Gel Bead and Library Kit V3.1 for samples (P/N 10000268, 1000120, PN-1000215 10x Genomics). GEM-RT was performed in a C1000 Touch Thermal cycler with 96-Deep Well Reaction Module (Bio-Rad; P/N 1851197): 53°C for 45 min, 85°C for 5 min; finally held at 4°C. After RT, GEMs were broken and the single-strand cDNA was cleaned up with DynaBeads MyOne Silane Beads (Thermo Fisher Scientific; P/N 37002D). cDNA was amplified using the C1000 Touch Thermal cycler with 96-DeepWell Reaction Module: 98°C for 3 min; 12 cycles of 98°C for 15 s, 63°C for 20 s, and 72°C for 1 min; 72°C for 1 min; held at 4°C. GEMs were then broken and the single-stranded cDNAs were cleaned up with DynaBeads MyOne Silane Beads (Thermo Fisher Scientific; P/N 37002D). The cDNAs were PCR amplified, cleaned up with SPRIselect beads (SPRI P/N B23318), fragmented, end-repaired, A-tailed, and size-selected with SPRIselect beads. Indexed adapters were ligated and cleaned up with SPRIselect beads. The resulting DNA fragments were PCR amplified and size-selected with SPRIselect beads. The size distribution of the resulting libraries was monitored using a Fragment Analyzer (Agilent Technologies, Santa Clara, CA, USA) and the libraries were quantified using the KAPA Library quantification kit (Roche, Basel, Switzerland). The libraries were denatured with NaOH, neutralized with Tris-HCl, and diluted to 100 pM. Clustering and sequencing were performed on a NovaSeq 6000 (Illumina, San Diego, CA, USA) using the paired-end 28-90 nt protocol on one lane of a flow cell SP.

#### Single-cell RNA-seq Analysis

Raw count matrices were imported and converted into Seurat objects (v5.3; ^50^). Each object was annotated by condition, i.e. APP^NL-F/NL-F^ or Control. Mitochondrial and ribosomal gene percentages were calculated to identify and exclude low-quality or stressed cells. Filtering thresholds were set at 5% for mitochondrial content and 20% for ribosomal content. Further quality controls were performed through the visualization of *nFeature_RNA* and *nCount_RNA* distributions, followed by visual thresholding adapted to each condition. Cells with fewer than 200 different genes were filtered out. Doublets were detected using the DoubletFinder algorithm (v2.0.6) with automatic estimation of multiplet rates, optimization of the *pK* parameter, and correction for homotypic doublets. Only cells classified as *singlet* were retained. Both datasets were then normalized using the Seurat Tool kit.

Gene selection was performed in two steps. First, the 5,000 most variable genes were selected using Seurat’s variance stabilization method and the integration performed using Seurat. Then to identify the most informative genes for cell classification, a Random Forest model (Ranger package, v0.17.0) was trained on real and permuted (simulated) cells using gene expression from the non-integrated objects to avoid model overfitting ^51^, finally leading to a list of 4,106 genes used for clustering.

Principal component analysis (PCA) was performed on the integrated data, and the top 22 principal components-accounting for 85% of the total variance - were retained for clustering. Clustering was performed using *FindClusters* with a resolution of 0.4. Of note, the stability of cluster assignments across different resolutions were assessed using the *Clustree* package (v0.5.1). Cluster annotation was initially carried out using SingleR (v2.8.0) and was further refined manually. Cluster proportions were determined across clusters and experimental conditions.

To identify dysregulated biological pathways between different clusters in WT conditions, or between control and AD conditions, we performed a Single Cell Pathway Analysis (SCPA) ^32^. Differentially expressed genes from microglial cells were used as input for enrichment against the Hallmark gene set collection (MSigDB v7.5.1; https://www.gsea-msigdb.org/). Pathway activity scores were computed at the single-cell level, allowing the comparison of pathway activation patterns between conditions. Statistical significance was assessed using permutation-based testing with false discovery rate (FDR) correction, and FDR < 0.05 considered significant. Results are shown as Q values, with Q=√−log10(padj).

## Results

### Early upregulation of Aβ42 in the APP^NL-F/NL-F^ mouse AD model

In their initial 2014 development of the APP^NL-F/NL-F^ model, Saito et al. investigated Aβ40 and Aβ42 expression from young adults (2-mo) to aged (24-mo) APP^NL-F/NL-F^ mice, showing that Aβ42 levels increased from 12/15-months of age ^12^. To characterize AD-related Aβ species during the earlier stages of amyloid pathology in APP^NL-F^ mice in maximum detail, we here used ultrasensitive ELISA-MSD technology to quantify cortical levels of Aβ38, Aβ40, and Aβ42 in both male and female homozygous mutant APP^NL-F/NL-F^ and APP^NL-F/WT^ heterozygous mouse cortex at ages from 3-12 months. We analysed both Triton X-100 (Tx-100)-soluble species, representing monomers, small oligomers and protofibrils, as well as guanidine (Gua)-soluble species, corresponding to aggregated fibrils ^13^. Small Aβ42 species were reliably detected in the Tx-100 soluble fractions of APP^NL-F/NL-F^ as early as 3 months of age, and in APP^NL-F/WT^ from 9 months onward. In the homozygote, Tx-100 soluble Aβ42 levels remained stable until 6 months, then increased in an age- and genotype-dependent, but sex-independent manner (Fig. 1A, for statistics see Supp-Table1).

**Figure 1:**
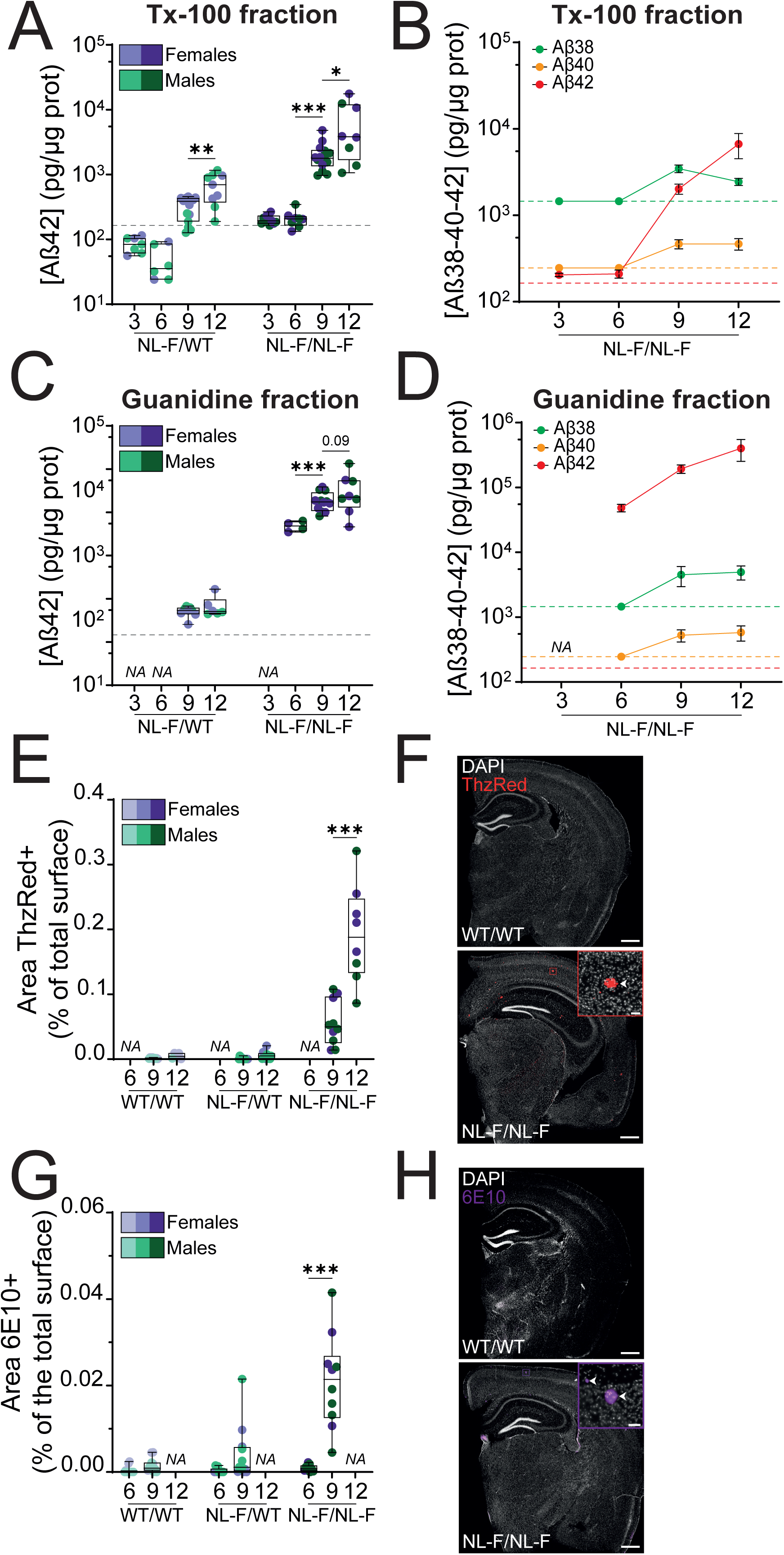
Amyloid species accumulate early in a genotype-dependant manner in the cortex of APP^WT/NL-F^ and APP^NL-F/NL-F^ mice. (**A**, **D**) ELISA-MSD quantification of Aβ species in cortical extracts from APP^NL-F^ mice aged 3, 6, 9 and 12 months. Tx-100-soluble fractions containing monomers, small oligomers and protofibrils are shown in (**A, B**) and guanidine-soluble aggregated fractions in (**C, D**). (**A, C**) quantification of Aβ42 in extracts of APP^WT/NL-F^ and APP^NL-F/NL-F^. (**B**, **D**) quantification of Aβ38, Aβ40 and Aβ42 in extracts of APP^NL-F/NL-F^ across ages. Aβ species were undetectable in samples from APP^WT/WT^ mice and are therefore not shown. The horizontal dotted line indicates ELISA-MSD detection limit (1456, 246 and 164 pg/µg for Aβ38, Aβ40, and Aβ42 respectively). In (**B, D**) data points located on the dotted lines represent values at or below the detection limit in the processed samples. (**E**, **G**) Quantification of Thiazine Red (ThzR)-positive mature amyloid beta, deposited plaques (**E**) and 6E10-positive formative diffuse plaque areas (**G**) in the cortex of APP^NL-F^ mice aged 3, 6, 9 and 12 months. (**F**, **H**) Representative images of ThzR staining in 12-mo mice (**F, red**) and 6E10 immunostaining in 9-mo mice (**H, purple**) from APP^WT/WT^ and APP^NL-F/NL-F^ genotypes. Arrows heads indicate amyloid deposits. Scale bar: 500 µm (insert: 50µm). Data are presented as box-and-whisker plots showing the median, interquartile range (25^th^–75^th^ percentiles), and minimum and maximum values (A, C, E, G) or as mean ± SEM (B, D). Each data point represents one animal. Sex is indicated by color (males, green; females, purple), while genotype is indicated by color shade (APP^WT/WT^, light; APP^NL-F/WT^, medium; APP^NL-F/NL-F^, dark). <u>Statistical analyses on the figure</u>: Mann-Whitney one-tailed between consecutive age groups within the same genotype. * p<0.05, ** p<0.01, *** p<0.001. “NA”: Not Analyzed. Equivalent quantification of ThzR- and 6E10-positive area in the hippocampus are shown in Supp-Fig. 1.

In contrast, Aβ38 and Aβ40 species were only detected later at later stages (9 and 12 months) and only in homozygous APP^NL-F/NL-F^. In this genotype, Tx-100-soluble Aβ42 levels were approximately fivefold higher than those of Aβ40. Notably, Aβ38 exceeded Aβ42 at 9 months, but this reversed by 12 months as Aβ42 increased and Aβ38 decreased (Fig. 1B). Because of the marked genotype and age-dependent differences, we analysed Tx-100 soluble Aβ42 levels separately in APP^NL-F/NL-F^ and APP^NL-F/WT^ mice, with successive age groups compared using one-tailed Mann-Whitney tests. In both homo- and hetero-zygotes, Aβ42 levels increased between 6- and 9-mo (APP^NL-F/NL-F^: p<0.001); and between 9- and 12-mo (APP^NL-F/WT^: p<0.01; APP^NL-F/NL-F^: p<0.05; Fig. 1A, Supp-Table1).

We also assessed aggregated guanidine-soluble fractions (Fig. 1C). These analyses highlighted aggregated Aβ42 as the predominant species in these fractions, with Aβ38 and Aβ40 barely detected (Fig. 1D). Although Aβ42 was detected in the heterozygote, its level in APP^NL-F/NL-F^ was about 400-fold higher levels compared to APP^WT/NL-F^. Similar to Tx-100 soluble fractions, no sex differences were observed in the global analysis (Fig. 1C, Supp-Table1). In sex-consolidated APP^NL-F/NL-F^ data, we observed a trend toward increased Aβ42 levels with age (1-way ANOVA – Age: p=0.07, F(2,21)=2.97). Successive age groups comparisons revealed a significant increase in aggregated Aβ42 in the homozygote between 6- and 9-mo (one-tailed Mann-Whitney p<0.001; Fig. 1C; Supp-Table1) with a further increase between 9- and 12-mo (APP^NL-F/NL-F^: p=0.09; Fig. 1C; Supp-Table1).

Complementing the above biochemical approach, we assessed cortical and hippocampal Aβ deposition histologically, in 9- and 12-mo mice with Thiazin-Red (ThzRed) stain. In addition to the cortex, hippocampus was examined, because it has been shown to be affected in APP^NL-F^ mice, albeit to a lesser extent than the cortex ^12^, and because of its critical role in memory processes. ThzRed specifically binds to β-sheet-rich structures in fibrillar amyloid aggregates ^14,15^, thus primarily detecting mature amyloid plaques. Consistent with our MSD results, no ThzRed-positive plaques were observed in either APP^WT/WT^ or APP^NL-F/WT^ mice. In contrast, ThzRed-positive amyloid plaques were consistently observed in APP^NL-F/NL-F^ mice starting at 9-months of age (Fig. 1E-F). The burden of cortical Aβ plaques increased in an age- and genotype-dependent but sex-independent manner (Fig. 1E; Supp-Table1). Aβ plaques were less abundant in the hippocampus than in the cortex; nevertheless, plaque burden in the hippocampus did increase over time, most strongly in homozygotes (Supp-Fig. 1A-B; Supp-Table1). Analyses of sex-consolidated data in both brain regions confirmed significant increase in the plaque burden between 9 and 12 months of age (Fig. 1E, Supp-Fig. 1A, Supp-Table1). These data are thus the earliest molecular phenotype of the APP^NL-F/NL-F^ mouse AD model; wherein mature brain plaques appear in the first year of life.

To investigate whether plaques could be detected at an even earlier age, we performed 6E10 immunostaining, which also detects formative diffuse plaques, in 6- and 9-mo mice. In the cortex, our results show an age- and genotype-dependent, but sex-independent 6E10 staining increase (Fig. 1G; Supp-Table1). Pairwise comparison further revealed a trend toward higher 6E10 immunostaining in 6-mo APP^NL-F/NL-F^ homozygotes mice compared to wild type (Mann-Whitney, p=0.0257; Supp-Table1), a difference that was not observed in the hippocampus (Supp-Fig. 1C; Supp-Table1). Thus, even at 6-months of age, slight brain differences can be observed in the APP^NL-F^ model. However, 9 months emerges as a pivotal timepoint at which amyloid burden significantly escalates in the homozygote. In contrast, only low amyloid burden and negligible dense amyloid plaques were observed in heterozygotes APP^NL-F/WT^ mice up to 12-months of age. We further show that both Tx100- and Guanidine-soluble Aβ42 levels, the predominant Aβ species in APP^NL-F/NL-F^, increase with age in both sexes of the homozygous AD mouse model.

### Aβ metabolism shifts without Tau alterations at 9-months of age in APP^NL-F/NL-F^

Given the increase in amyloid burden at 9 months in homozygous APP^NL-F/NL-F^ mice, we examined whether this stage was marked by alterations in various markers of AD progression i.e. APP expression, Aβ-metabolizing enzymes, or Tau-related pathology. We first validated the stable expression of the human APP protein in the APP^NL-F/NL-F^ mice (Supp-Fig. 2A). We then examined whether the increased amyloid load was associated with alterations in Aβ metabolism, by quantifying the expression of IDE (Insulin-Degrading Enzyme) and BACE1 (β-secretase1), two key enzymes involved in Aβ production and clearance. IDE expression was unaffected by sex or mutation, though male APP^NL-F/NL-F^ displayed marked variability, with some exhibiting higher expression (Supp-Fig. 2B). In contrast to the sex-independent increase in amyloid burden, BACE1 expression exhibited clear sex- and genotype-specific differences. In males, BACE1 levels were significantly reduced in mutation carriers as compared to APP^WT/WT^ mice, while APP^NL-F/NL-F^ and APP^WT/WT^ females had little BACE1expression (Supp-Fig. 2C).

Although amyloid mouse models typically do not develop overt Tau pathology ^16^, Aβ deposition has been implicated in promoting Tau phosphorylation ^17^, a key pathological modification which contributes to Tau aggregation and the formation of neurofibrillary tangles ^18^. To evaluate potential Tau-related changes, we assessed cortical Tau phosphorylation in 9-mo mice by Western blot using the AT8 antibody (detecting p-S202/p-T205 epitopes). At this disease stage, however, we did not detect significant changes in either total Tau, as assessed using AT5 antibody, nor in phospho-Tau/total Tau ratios (Supp-Fig. 2D-E). Overall then, these data indicated that amyloid accumulates at 9 months of age in the APP^NL-F/NL-F^ model, without detectable changes in IDE or Tau phosphorylation, but there is BACE1modulation in males. This indicates that, at this stage, increased amyloid burden is not accompanied by broad and consistent alterations in Aβ-clearing pathways or Tau-related pathology.

### Microgliosis and astrogliosis are detected from 9-months old in APP^NL-F^ homozygotes

Microgliosis and astrogliosis are key hallmarks of Alzheimer’s disease. To characterize the temporal dynamics of these pathological features at early stages in APP^NL-F^ mice, we used immunohistochemistry to quantify IBA1 (microglia) and GFAP (astrocyte)-positive areas in the cortex and hippocampus at 6-, 9-, and 12-months of age across the three genotypes, while also assessing any potential sex differences. There was no evidence of microgliosis or astrogliosis at any time point in the cortex or hippocampus of APP^WT/WT^ or APP^NL-F/WT^ heterozygotes (Fig. 2A,C; Supp-Fig. 3A,C). In contrast, homozygous APP^NL-F/NL-F^ mice exhibited microglial and astrocytic clustering around amyloid plaques as shown by co-staining of IBA1, GFAP and ThzRed respectively (Fig. 2E).

**Figure 2:**
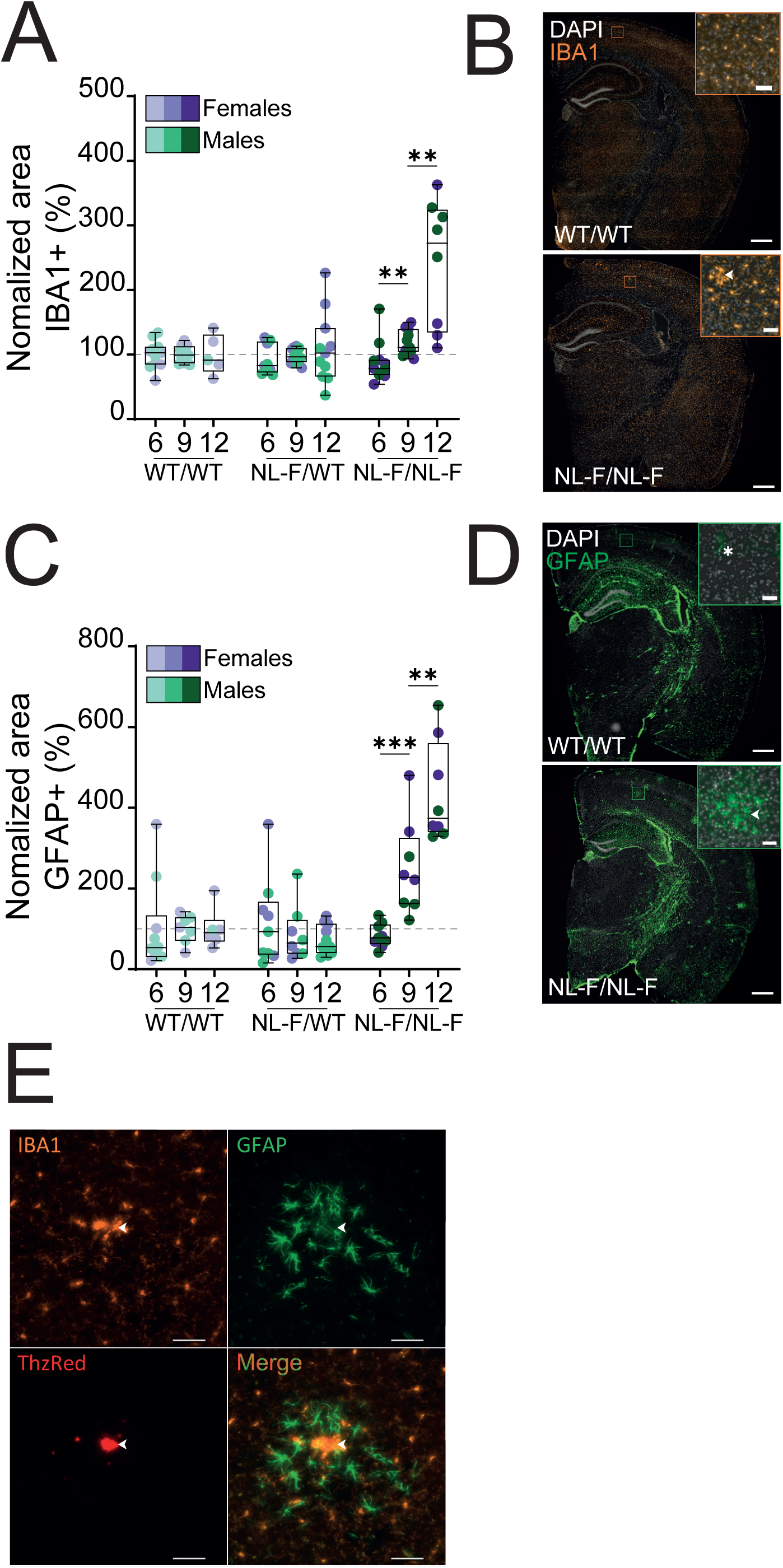
Progressive increase in microgliosis and astrogliosis in the cortex of APP^NL-F/NL-F^ mice during the development of amyloidosis. (**A**) Quantification of cortical-IBA1 (microglia) positive area in 6-, 9- and 12-month-old APP^NL-F^ mice, normalized to aged matched controls. (**B**) Representative IBA1 immunostaining of 12-mo APP^WT/WT^ and APP^NL-F/NL-F^ mice. Arrow head in insert show microglia clustering. Scale bar: 500µm; insert: 50µm. (**C**) Quantification of cortical-GFAP (astrocyte) positive area in 6-, 9-and 12-month-old APP^NL-F^ mice, normalized to aged matched controls.(**D**) Representative GFAP immunostaining of 12-mo APP^WT/WT^ and APP^NL-F/NL-F^ mice. Star and arrow head in insert shows vessels-associated astrocytes (APP^WT/WT^) and astrocyte clustering (APP^NL-F/NL-F^) res pectively. Scale bar: 500µm; insert: 50µm. (**E**) Representative immunostaining of microglia (IBA1) and astrocyte (GFAP) clustering around amyloid plaque (ThzRed) in the cortex of a 12-mo APP^NL-F/NL-F^ mice. Scale bar: 500μm; insert: 50μm. Merged area show a merge of IBA1, GFAP and ThzRed staining. Quantitative data are shown as box-and-whisker plots indicating the median, interquartile range (25^th^–75^th^ percentiles), and minimum and maximum values. Each data point represents one animal. Sex is indicated by color (males, green; females, purple), while genotype is indicated by color shade (APP^WT/WT^, light; APP ^NL-F/NL-F^, dark). Horizontal dotted line represents 100% of the mean control value. <u>Statistical analyses on the figure</u>: one-tailed Mann-Whitney between consecutive ages within a same genotype. ** p<0.01, *** p<0.001. Equivalent data for hippocampus is shown in Supp-Fig. 3.

Quantification revealed significant age-dependent microgliosis in both cortex and hippocampus, with no significant difference between males and females (Fig. 2A; Supp-Fig. 3A; Supp-Table1). Analysis of sex-consolidated data using one-tailed Mann-Whitney tests showed a significant increase in IBA1+ area in APP^NL-F/NL-F^ mice between 6 and 9 months in both the cortex (p<0.01; Fig. 2A; Supp-Table1) and the hippocampus (p<0.001, Supp-Fig. 3A, Supp-Table1), with a further significant increase from 9 to 12 months (Cortex Fig. 2A & Hippocampus Supp-Fig. 3A; p<0.01; Supp-Table1).

In wild-type APP^WT/WT^ and heterozygous APP^NL-F/WT^ mice, GFAP+ astrocytes were sparsely distributed in the cortex, mostly along vessel-like structures (Fig. 2D, top panel), with no evidence of astrogliosis (Fig. 2C). Quantification revealed significant age-dependent astrogliosis in the cortex of APP^NL-F/NL-F^ homozygous mutants from 9-mo with no detectable sex difference (Fig. 2C; Supp-Table1). Mann-Whitney pairwise comparisons across successive age groups in homozygous mutants confirmed a significant increased astrogliosis from 9-months onward (Fig. 2C), whereas in hippocampus it was only detected at 12 months (Supp-Fig. 3C, p<0.001). Overall, our results demonstrate a significant age-dependent increase in both microgliosis and astrogliosis in the cortex and the hippocampus of both male and female homozygous APP^NL-F^ mice. In the cortex, both microgliosis and astrogliosis were evident from 9-months of age, whereas hippocampal astrogliosis emerged more gradually becoming prominent by 12 months. Added to the first detection of Aβ42 in the first section, these data represent the earliest cellular phenotype detected in this mouse model of AD.

### Cognitive deficits in 9-mo APP^NL-F^ homozygotes

In their initial characterization of the APP^NL-F/NL-F^ mouse model, Saito et al. ^12^ reported deficits in Y-maze spontaneous alternation beginning at 18 months of age. However, subsequent studies have identified behavioural deficits at earlier stages ^19–21^.

Although learning and memory deficits in AD are more closely associated with Tau pathology, mouse models of amyloidosis also exhibit cognitive impairments ^16^. Importantly, sensitive behavioural assays are required to detect early functional consequences of pathology ^22,23^.

To comprehensively assess early-stage cognitive alterations, we employed a validated behavioural assessment protocol. This included five days of experimenter handling to minimize stress, followed by open-field testing to assess locomotor activity and anxiety-like behaviour. We concluded with an eight-day Barnes Maze trial to evaluate the visuo-spatial learning and memory (Fig. 3A). To examine early deficits and potential sex differences, we evaluated 8–14 mice per genotype and sex, including 69 mice at 6 months and 61 at 9 months of age. Open-field analysis revealed older animals covered less distance over the 10 minutes of the test, but neither sex nor genotype significantly influenced locomotor activity (Supp-Fig. 4A-B; Supp-Table 1). Similarly, anxiety-like behaviour, measured as the percentage of time spent in the centre of the arena, was impacted by age, but unaffected by sex or genotype at either 6 or 9 months (Supp-Fig. 4C-D; Supp-Table 1).

**Figure 3:**
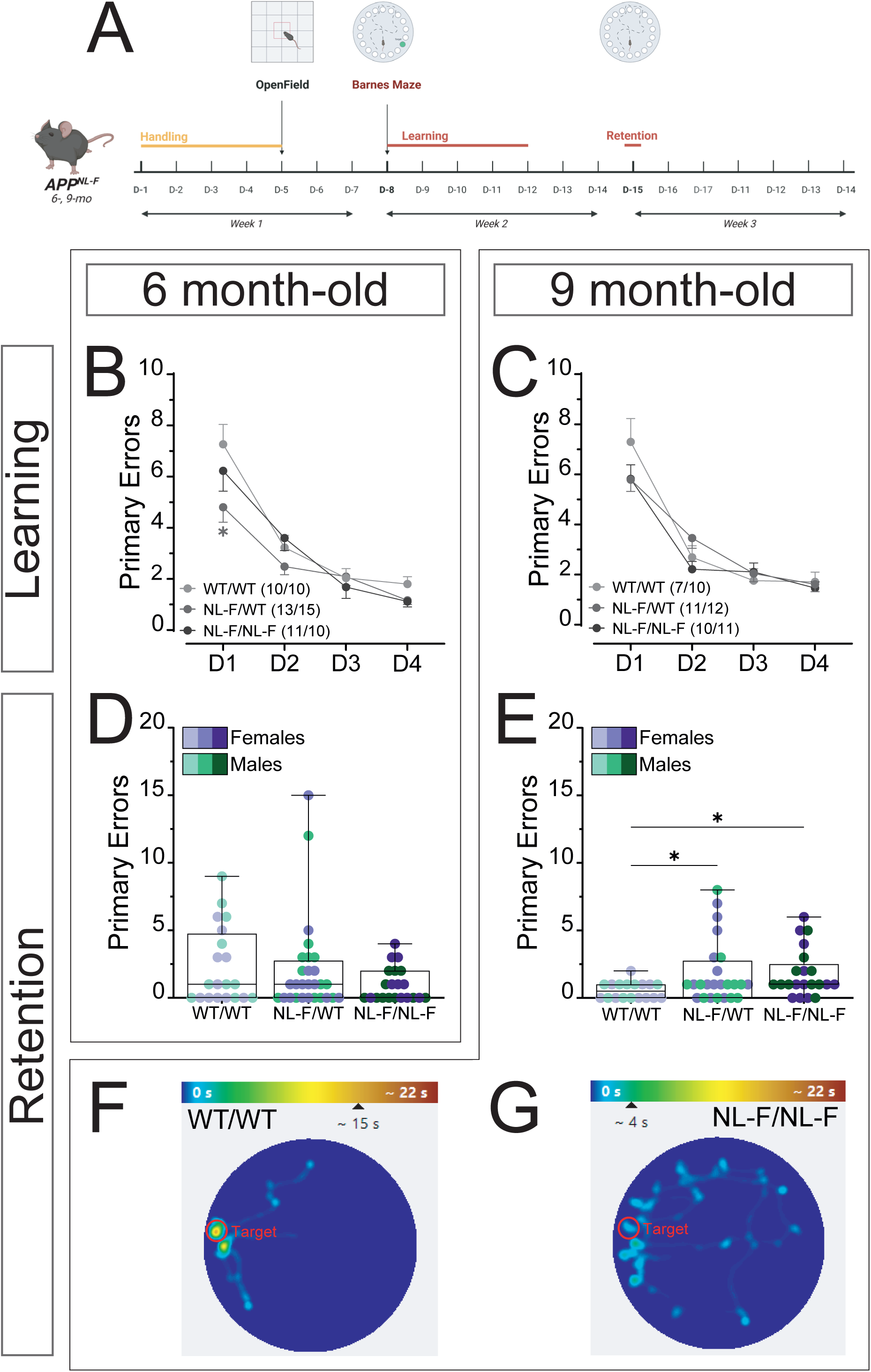
Progressive impairment of cognitive performances in APP^NL-F^ mice during amyloidosis. (**A**) Schematic of the behavioral protocol study used to assess cognitive impairment. (**B**, **C**) Learning performance, quantified as the number of errors, in 6-mo (**B**) and 9-mo (**C**) APP^NL-F^ mice during the Barnes maze over the 4-days learning phase. Numbers in parentheses indicate the number of male and female mice analysed in each group. (**D**, **E**) Retention performance of 6-mo (**D**) and 9-mo (**E**) APP^NL-F^ mice in the Barnes maze retention phase. (**F**, **G**) Representative traces and heatmaps of individual 9-mo APP^WT/WT^ (**F**) and APP^NL-F/NL^ ^F^ (**G**) mice during the retention task. Data are shown as mean ± SEM (**B**, **C**) or box-and-whisker plots showing the median, interquartile range (25th–75th percentiles), and minimum and maximum values (**D**, **E**). Each data point represents one animal. Sex is indicated by color (males, green; females, purple), while genotype is indicated by color shade (APP^WT/WT^, light; APP^NL-F/WT^, medium; APP^NL-F/NL-F^, dark). <u>Statistical analyses</u>: one-tailed Mann-Whitney test between genotypes mice within the same age group. * p<0.05.

In the absence of locomotor or anxiety defects we then tested learning ability in the Barnes Maze by quantifying primary errors during reaching of the target hole. Acquisition profiles were analysed separately at 6 and 9 months using a 3-way linear mixed-effects model (LMM) with sex and genotype as between-subject factors and training session as within-subjects repeated-measure factor. At 6 months, no significant sex effect was detected (Supp-Table 1), allowing data to be consolidated across sexes to increase statistical power for genotype comparisons. A genotype effect was observed on day 1, with APP^NL-F/WT^ mice making fewer errors than wild type controls (Fig. 3B, Supp-Fig. 4E and Supp-Table 1). However, all groups, including APP^NL-F/NL-F^, showed progressive reductions in primary errors across training sessions, indicating successful task acquisition. At 9 months, females made modestly but significantly more errors than males, but this effect was independent of genotype (Supp-Table1). Given the limited and genotype-independent effect of sex, data were pooled for subsequent genotype-focused analyses. During the acquisition phase, primary errors decreased over time in all genotypes (Fig. 3C, Supp-Table 1), indicating intact learning in all groups.

In the Barnes maze probe trial, which assesses spatial memory retrieval after task acquisition, all 6-mo mice performed better than during their final training session, indicating intact memory recall, with no deficit detected in either APP^NL-F/WT^ or APP^NL-F/NL-F^ mice (Fig. 3D; Supp-Fig. 4G). At 9 months of age, APP^WT/WT^ mice likewise showed improved performance relative to their final training session, consistent with preserved spatial memory. In contrast, despite having learned the task during training, both APP^NL-F/WT^ and APP^NL-F/NL-F^ exhibited impaired memory retrieval, making more errors than wild-types controls before locating the target hole (Fig. 3E; Supp-Fig. 4H). 9-month-old APP^NL-F/NL-F^ mice, relied less on direct spatial search strategies, with mice adopting more disorganized and inefficient exploration patterns to locate the target (Fig. 3F-G). Overall, our comprehensive analyses revealed small but significant deficits in learning and memory performances in 9-mo APP^NL-F^ mice, particularly in the homozygous APP^NL-F/NL-F^. This suggests that subtle cognitive alterations are detectable at earlier stages of AD-like pathology in this model than had previously been thought.

### Global gene deregulation in 9-mo APP^NL-F^ homozygotes

To explore the impact of genotype, age and sex on brain inflammation, we took a targeted qPCR approach to measure the expression levels of 22 genes known to be deregulated under neurodegenerative conditions and/or specifically in AD, in 6-, 9- and 12-mo male and female mouse cortex ^8,24^. We chose mostly microglia-specific genes and/or genes that are deregulated in reactive microglia (*Hexb*, *P2ry12*, *Ccl12*, *Cd68*, *Cst7*, *Clec7a*, *Trem2*, *Tyrobp*, *Lpl*, *Gpnmb*, *Itgax*, *Spp1* & *Ctss*). We also included several markers of astrocyte activation (*Gfap*, *H2T23*, *Tmsf4*, *Serpina3n*) plus a selection of neuroinflammation-related genes (*Cxcl10*, *Il1b*, *Il6*, *Nlrp3*, *Syk*). Gene expression was analysed at each stage of the pathology using 2-way ANOVA, with genotype and sex as fixed factors, as well as their interaction. The main results for 9-mo mice are presented in Figures 4 and 5, while complementary analyses at this age and the results obtained in 6- and 12-mo mice are shown in Supp-Fig. 5-7 (see also statistics in Supp-Table1). Principal component analysis (PCA) and heatmaps were used to explore the transcriptional variability among genotypes and sex. These preliminary analyses revealed distinct basal and genotype-dependant transcriptomic profiles in males and females, thus sexes were studied independently (Fig. 5A-B, Supp-Fig 6A,7A). At 6-months of age, neither PCA or heatmap analyses separated the samples from the different groups (Fig. 4A and Supp-Fig. 5A, Supp-Fig. 6A-B). The only gene that was noticeably affected by genotype was *Cst7* in females (Supp-Fig. 6F). In 9-mo males PC1 accounted for 35.5% of the variance, yet it did not clearly separate the groups (Fig. 4B). However, the second principal component PC2 accounted for 17.5% of variance and distinguished APP^NL-F/NL-F^ mice, from both APP^WT/WT^ and APP^NL-F/WT^. This molecular profile was driven by high contributions from inflammatory chemokine genes such as *Tyrobp* and *Ccl12* (Fig. 4B). In females, the first principal component, PC1, explained a larger proportion of the variance (42.5%) with separation of APP^NL-F/NL-F^ individuals from the other groups (Fig. 4B). Genes that drive the separation along PC1 included *Cst7*, *Clec7a*, *Trem2*, *Hexb* and *Syk* (Fig. 4B). The specific expression profiles of some of the genes with high contribution to the separation between the experimental groups at 9-months of age are shown in Fig. 5C-H. At this later timepoint *Cst7* was upregulated in both male and female APP^NL-F/NL-F^. The *Clec7a*, *Hexb* and *Trem2* genes were also up-regulated in female and *Tyrobp* in male mutant cortex.

**Figure 4:**
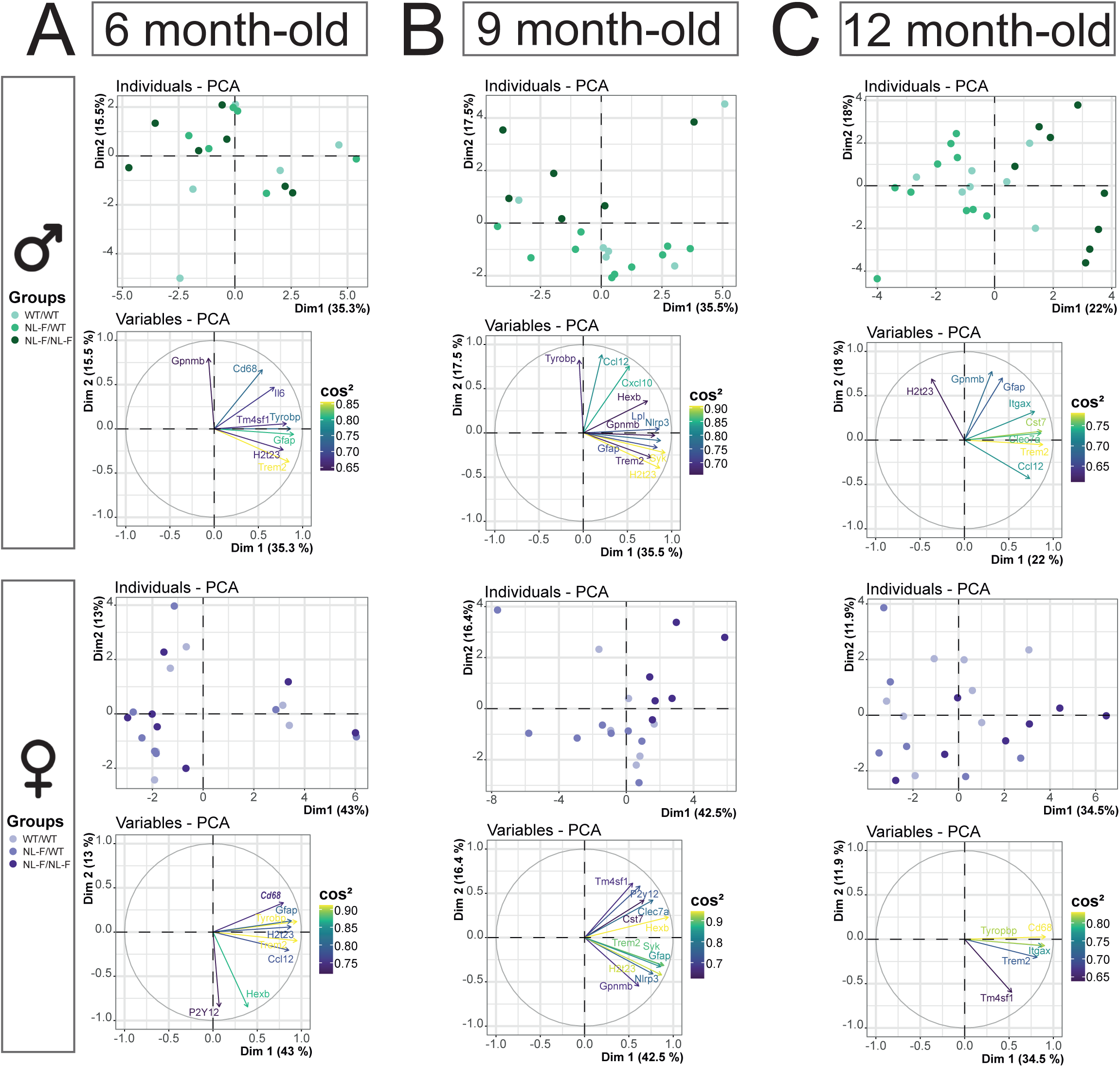
Gene expression changes in cortical homogenates from 6-, 9- and 12-mo APP^NL-F^ for a range of inflammation and AD-related genes. **(A-C)** PCA plots of individual samples and corresponding gene loading plots (cos² > 0.6) projected along PC1 and PC2 in 6- (**A**), 9- (**B**) and 12-mo (**C**) samples. Data are presented separately for males (top panels) and females (bottom panels). Each data point represents one animal. Sex is indicated by color (males, green; females, purple), while genotype is indicated by color shade (APP^WT/WT^, light; APP^NL-F/WT^, medium; APP^NL-F/NL-F^, dark). Scaled component value (cos²) are displayed using a purple–yellow color scale, with yellow indicating higher representation of a given gene on the component and purple indicating lower representation.

**Figure 5:**
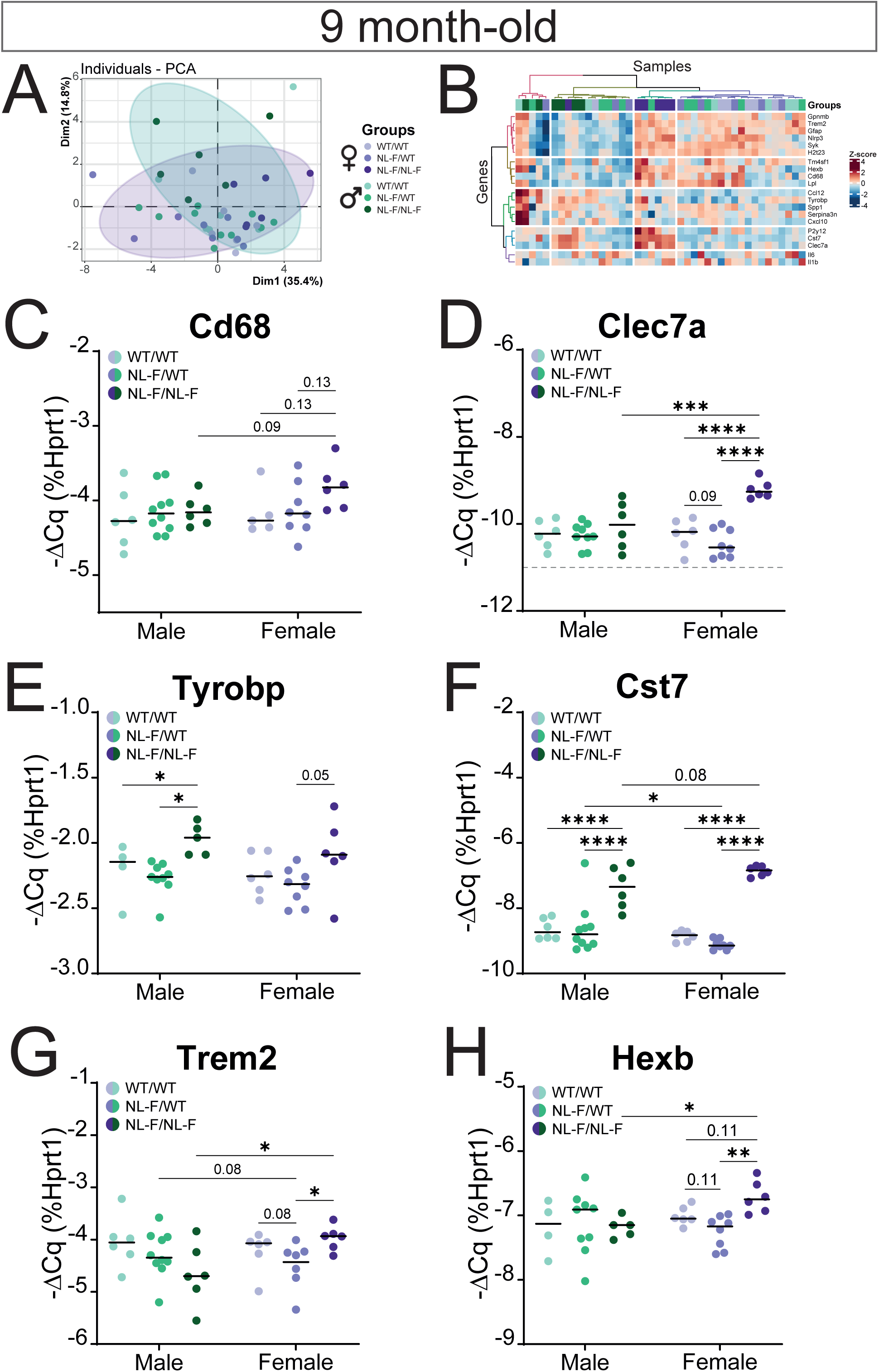
Gene expression changes in the 9-mo mouse cortex homogenates for a range of inflammation and AD-related gens. Related to Figure 4. (**A**) PCA plot of individual samples projected along PC1 and PC2. (**B**) Heatmap of differentially expressed genes. Scaled expression values (row Z score) are shown using a blue–red color scale with red indicating higher expression, and blue lower expression. (**C**-**G**) Gene expression changes measured by qPCR and plotted as −ΔCq for a selection of genes. Data are shown as dot plots indicating the median. Each data point represents one animal. Sex is indicated by color (males, green; females, purple), while genotype is indicated by color shade (APP^WT/WT^, light; APP^NL-F/WT^, medium; APP^NL-F/NL-F^, dark). <u>Statistical analysis for each selected gene</u>: 2-way ANOVA with genotype and sex as between subjects’ factors. FDR corrected post-hoc tests. * p<0.05.

In 12-mo males, the PC1 clearly separated APP^NL-F/NL-F^ homozygotes from both APP^WT/WT^ and APP^NL-F/WT^ along PC1, indicating significant transcriptional changes emerge only in homozygous mutants (Fig. 4C). In contrast, APP^NL-F/WT^ and APP^WT/WT^ individuals overlap substantially, indicating that heterozygous mutations do not strongly alter transcriptomic profiles at this age. Genes that drive the separation along PC1 were *Cst*7, *Trem2*, *Clec7a*, and to a lesser extend *Itgax* and *Ccl12* (Fig. 4C). Specific expression profiles of the most discriminative genes are shown in Supp-Fig. 7C-H. In 12-mo female, we also observed a trend for a separation of the APP^NL-F/NL-F^ mice along PC1, but with substantial variability within this group, with notably three mice showing transcriptomic profiles that globally resembled that of control or heterozygous mice (Fig. 4C). However, all genes analysed individually (i.e. *Cd68*, *Clec7a*, *Cst7*, *Trem2* and *Tyrobp*) were significantly up-regulated in 12-mo female APP^NL-F/NL-F^ mice compared to controls (Supp-Fig. 7C-H). Notably, *Cst7* and *Clec7a* remained the most upregulated genes in both males and females. Overall, our targeted qPCR analysis, revealed that, unlike the heterozygotes, APP^NL-F/NL-F^ homozygous mice exhibit age- and sex-dependent alteration in neuroinflammatory genes increasing from 9 months.

### An early global microglial transcriptional shift in the APP^NL-F/NL-F^ amyloid pathology model

Our targeted qPCR approach thus pointed to early microglia dysregulation in the cortex of the APP^NL-F/NL-F^ AD model mouse. To further delineate microglia reaction in this amyloid pathology model, we performed single-cell RNA-seq (scRNA-seq) on cortical brain myeloid cells isolated from 12-mo APP^NL-F/NL-F^ mice and littermate controls. The 12-month timepoint was chosen to characterize the established microglial response once amyloid pathology was well established, allowing for a clear definition of the cellular states associated with a chronic amyloid environment. Briefly, after mechanical and enzymatic dissociation, myeloid cells were isolated based on membrane expression of CD11b using magnetic-activated cell sorting (MACS; Fig. 6A). Preliminary experiments revealed that this approach allows the isolation of brain macrophages, with microglia, as expected, representing over 90% of the CD11b+ cells ^25^. We then mapped the cellular heterogeneity of brain myeloid cells in 12-mo mouse brain and analysed the specific changes associated with early-stage amyloidosis, by performing droplet-based scRNA-seq using 10X Genomics Chromium technology. Overall, 10,590 cells (i.e. 5,860 cells APP^WT/WT^ controls and 4,730 cells from APP^NL-F/NL-F^ mice; Fig. 6B) passed quality control controls and were included in the dataset. To delineate the clusters, we used the Clustree package ^26^ to assess the evolution of hierarchical clustering among all cells, hinting at aberrant sub-clustering. Twelve different clusters, annotated C0 to C11, from the largest to the smallest in terms of cell number, were identified (Fig. 6B,C).

**Figure 6:**
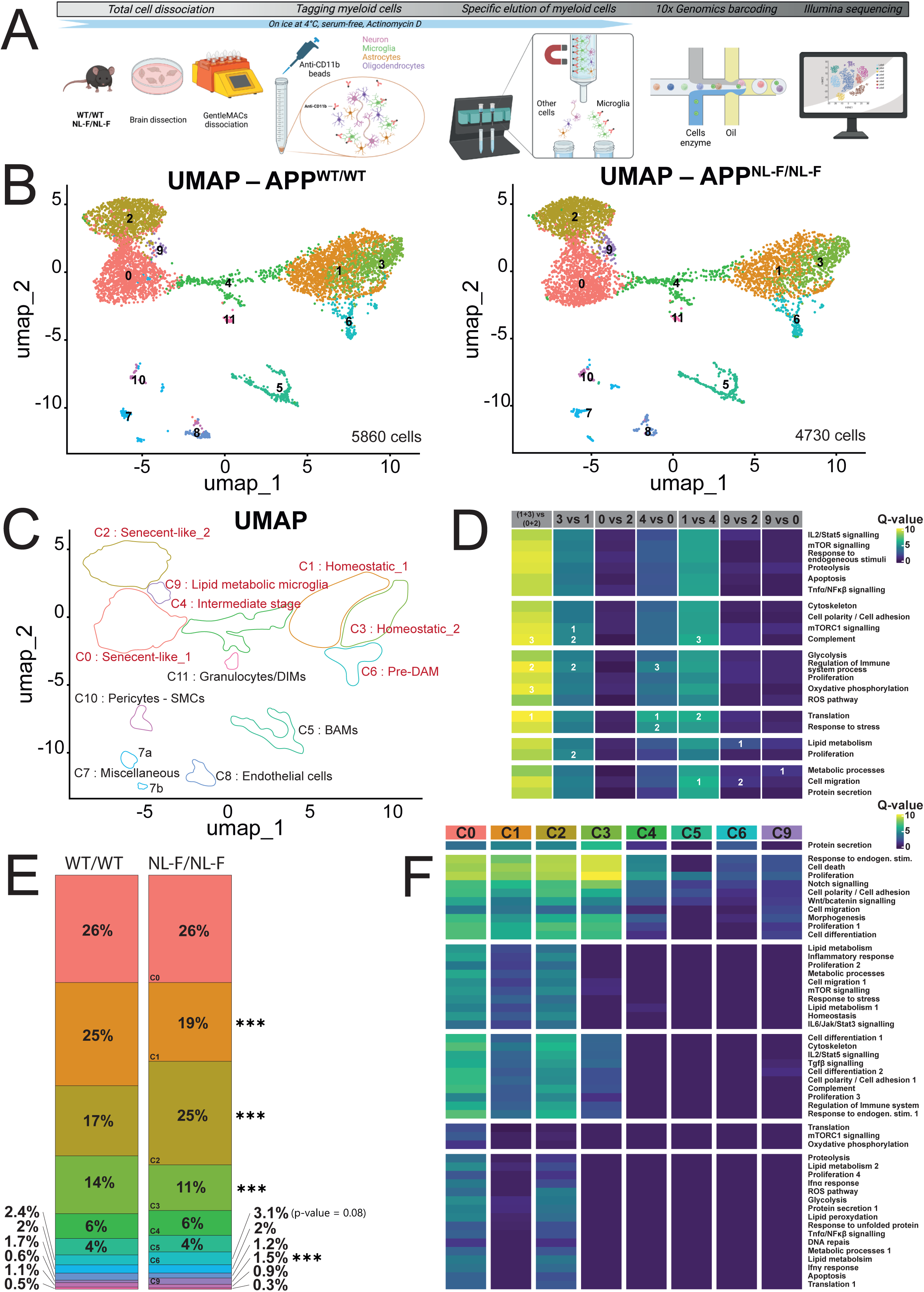
Single-cell transcriptomic landscape of cortical CD11b⁺ cells in APP^NL-F/NL-F^ mice. (**A**) Schematic overview of the cell isolation and scRNA-seq workflow (**B**) UMAP plots showing the twelve clusters of cortical CD11b^+^cells from scRNA-seq data obtained in APP^WT/WT^ (left panel) and APP^NL-F/NL^ ^F^ (right panel) mice. (**C**) Schematic summarizing cluster identities and nomenclature. (**D**) Heatmap of Q values generated by single-cell pathway (SCPA) analysis. The union of Top-10 Hallmark pathways identified in each comparison is shown, with top three pathways per comparison indicated. (**E**) Relative proportion of cells within each cluster in APP^WT/WT^ and APP^NL-F/NL^ ^F^ conditions. (**F**) Heatmap of Q values from SCPA comparing APP^WT/WT^ and APP^NL^ ^F/NL-F^ cells within each cluster. All Hallmark pathways are shown.

The SingleR package was first used for automatic identification of cell types based on a previous mouse RNA-seq annotation ^27^. As expected from the isolation strategy, seven of the twelve clusters (i.e. C0, C1, C2, C3, C4, C6 and C9) were identified as microglial cells, based on the expression of canonical microglial markers such as *Pr2y12*; *Sall1*; *Hexb* and *Tmem119* (Supp-Fig. 8A,B). Cluster 5 was identified as Border Associated Macrophages (BAMs) based on their expression of typical BAM markers, including *Mrc1* and *F13a1* (Supp-Fig. 8A,C). Minor Clusters 7 and 8 each represented approximatively 2% of the total cell population. Cluster 7 was a mixed cell population composed of neurons, astrocytes, and oligodendrocytes (Supp-Fig. 8A,D,E). Indeed, within this cluster, the larger subcluster (7a) displayed enriched expression of *Dclk1* (Supp-Fig. 8A, neuronal marker) but also *Gfap*, *Fgfr3*, *Gja1* and *Aqp4* (Supp-Fig. 8D, astrocyte markers) whereas the smallest (7b) was characterized by higher expression of *Plp1*, *Mog* and *Mobp* (Supp-Fig. 8E, oligodendrocytes markers). Cluster 8 was identified as endothelial cells and was characterized by increased expression of *Flt1*, *Jcad* as well as *Cldn5*, *Cdh5*, *Pecam1* and *Vwf* (Supp-Fig. 8A,F). Cluster 10 and 11 were sparse, representing respectively only about 1% and 0.5% of the total number of cells, and comprising just over 100 cells and fewer than 100 cells, respectively. Of note, Cluster 11 was identified as correlated with the granulocyte signature and may represent the Disease Inflammatory Macrophages (DIMs) identified by Silvin et al. ^28^, exhibiting high expression of some DIM markers, including *S100A9* and *Slpi* (Supp-Fig. 8A,G).

Among the C0-C4, C6, C9 microglial clusters, Clusters 0, 1, 2, 3 were the most abundant microglial clusters. Although all clusters were microglial in identity, C0 closely resembled C2, while C1 showed strong similarity to C3. Clusters 1 and 3, which represent around 40% of all the cells, had higher expression of typical homeostatic microglia genes (including *Hexb*, *P2ry12*, *Sall1*, and *Tmem119*) than C0 and C2 (Supp-Fig. 8B). Consistent with this observation, C1/C3 exhibited higher expression of *Olfml3* (olfactomedin-like 3) which encodes a secreted extracellular matrix protein whose expression is directly regulated by TGFβ1/Smad2 signaling ^29^, an essential pathway for homeostatic state maintenance ^30^ (Supp-Fig. 8A). The C1/C3 clusters also expressed more *Rgs2* (Regulator of G-protein Signaling 2), a GTPase-activating protein that modulates GPCR signaling, and which has been shown to dampen TLR4-driven inflammatory signaling in innate immune cells ^31^.

To gain further insight into the functional difference between these distinct pairs of microglial states (i.e. C0+C2 vs. C1+C3), we performed single cell pathway analysis on APP^WT/WT^ control cells using the SCPA R package ^32^. We focused on the Hallmark collection of the MSigDB (Molecular Signatures Data Base) that represents well defined biological states. Comparing clusters C1+C3 and C0+C2 (Fig. 6D), SCPA analysis identified Translation, Regulation of Immune processes, Complement and Oxidative Phosphorylation as the top fundamental biological processes enriched in Clusters C1/C3 (Fig. 6D and Supp-Table 2). Overall, these results suggested that C1/C3 microglia exhibit a balanced, surveillance-ready phenotype, i.e. these are homeostatic microglia that survey the parenchyma and are ready to promptly respond to environmental perturbations. We thus refer to C3 and C1 as Homeostatic microglia respectively (Fig. 6C). Of note, compared to C1-, C3-microglia exhibited enrichment in pathways associated with cell polarity, regulation of immune processes, mTOR signaling, complement system and cell cycle, suggesting that C1-microglia may exhibit a pre-activated state, consistent with increased expression of genes associated with lysosomal (*Cd63*, *Ctsd*, *Cd68*), mitochondrial (*Tomm20*, *Ucp2*) and oxidative stress (*Hmox*, *Sod2*) (Supp-Fig. 9A-E). Single-cell pathway analysis did not identify any enriched pathway in the C0/C2 cluster over C1/C3. However, in addition to the reduced expression of canonical homeostatic microglial markers described above, the C0+C2 cluster, which together represent about 40% of the cells in APP^WT/WT^ cells– exhibited increased expression of *Malat1* (Supp-Fig. 8A & Supp-Fig. 9H), a long noncoding RNA known to be upregulated in immune cell subtypes in association with aging and frailty ^33^. This transcriptional profile, together with elevated expression of *Lysmd4*, *Mef2a* and *Tgfbr1*, suggests that the C0 and C2 clusters may represent microglia engaged in an aging/senescent-associated program (Supp-Fig. 9I-K). Notably, SCPA analysis did not identify any significantly enriched pathway in C0 *vs* C2 or vice versa (Fig. 6D, Supp-Table-2), supporting the close similarity between these two clusters. C0 and C2 were thus referred to as senescent-like microglia (Fig. 6C).

C4 microglia made up about 6% of the cells and seemed to represent a transitional state between C0 and C1, showing enrichment in biological functions related to translation, response to stress, and regulation of Immune processes (Fig. 6D, Supp-Table 2). Compared to C1, C4 exhibit less migration, translation, and complement related pathways (Fig. 6D, Supp-Table 2). C6-microglia represented 2.5% of the APP^WT/WT^ cells and selectively expressed elevated levels of the typical DAM markers, including *Lpl*, *Itgax*, *Clec7a* and *Cst7* (Supp-Fig. 9L-O). Of note C6 also exhibited high level of canonical microglial homeostatic genes (Supp-Fig. 8A), suggesting it does not a represent *bonafide* DAM microglia state. We thus refer to C6 as a pre-DAM cluster (Fig. 6C). C9-microglia represented only 0.6% of the APP^WT/WT^ cells. Compared to C0- and C2-microglia, C9-microglia were enriched in genes related to lipid metabolism, metabolic processes, and cell migration (Fig. 6D, Supp-Table 2). C9 also shows high expression of *ApoE* (Supp-Fig 8A) and C9 was thus referred to as Lipid-metabolic microglia (Fig. 6C).

Finally, we investigated how amyloid pathology influenced the relative proportions of the different cell clusters. In early-stage amyloidosis, in homozygous mutants we observed a significant reduction in the frequency of both C1 and C3 homeostatic microglia, whereas the proportion of C2 senescent-like microglia increased by nearly 50% suggesting amyloid pathology causes a shift in microglial composition from homeostatic (C1/C3) to senescent-like (C2) profiles (Fig. 6E). Interestingly, in APP^NL-F/NL-F^ mice, the relative proportions of C9 lipid-metabolic microglia and, to a lesser extent, C6 pre-DAM microglia also increased, by approximately 150% (p<0.001) and 30% (p=0.08), respectively. Notably, the relative proportion of C5-BAM was not affected at this early stage of the pathology.

To further investigate how amyloid pathology alters transcriptomic programs of different microglia/macrophage subtypes (i.e. C0-5, C6 and C9), we performed SCPA analysis ^32^, comparing APP^NL-F/NL-F^ and APP^WT/WT^ cells within each cluster using the HALLMARK gene set collection as reference (Fig. 6F, Supp-Table 3). Amyloid pathology had limited impact on HALLMARK gene sets in C5-BAM, C6-preDAM and C9-Lipid-metabolic microglia. In contrast, SCPA identified a substantially larger number of amyloidosis-enriched HALLMARK pathways for C1 and C3-homeostatic microglia, and an even greater enrichment in C0 and C2-sensescent-like microglia (Fig. 6F). Positive and negative responses to endogenous stimuli, cell proliferation and cell death were particularly enriched in APP^NL-F/NL-F^ microglia from all four, homeostatic- and senescent-like clusters (Fig. 6F, Supp-Table3). These cells also likely experience morphological changes (Cell polarity/cell adhesion; Morphogenesis) during amyloidosis. In addition to these alterations, APP^NL-F/NL-F^ C0- and to a lesser extent C2-senescent microglia were also enriched in inflammation (i.e.: Inflammatory response, IL6/Jak/Stat3 signalling, IL2/Stat5 signalling, Ros pathway) and lipid metabolism related pathways. Notably, these two microglial states were also enriched for pathways related to the response to unfolded protein, consistent with cellular stress responses associated with amyloid accumulation (Fig. 6F, Supp-Table3).

Overall, these analyses revealed that amyloidosis perturbs the transcriptomic profiles of the major microglial cell states. Senescent-like microglia increase in proportion in APP^NL-F/NL-F^ and exhibit the most altered transcriptional programs, whereas homeostatic-like microglia decrease in frequency and are also transcriptionally disturbed in the APP^NL-F/NL-F^ mice.

## Discussion

Alzheimer Disease is a progressive and irreversible disease and a major public health issue in the aging population. Pathological processes linked to amyloid deposition and Tau alterations begin decades before appearance of the clinical symptoms ^34^, underscoring the need for animal models that recapitulate the earliest stages of disease progression. As a second-generation AD mouse model, the APP^NL-F^ mouse offers several advantages including progressive amyloidosis that emerges around midlife. While this model has been characterized by multiple groups, most prior studies focused on amyloidosis and behavioural outcomes at limited or widely spaced time points [30, 33, 34, 45, 8, 25, 32], leaving the early-stage dynamics of pathology poorly defined. Here, we extend these observations by providing a detailed longitudinal analysis from 3- to 12-months of age, with a particular emphasis on early neuroinflammatory processes. Our results highlight a key interaction between amyloid pathology and microglial aging as a potential driver of disease progression.

By systematically charting molecular, histological, and functional alterations, we identify a pivotal inflection point around 9-months of age, marking the transition from a pre-symptomatic to an early pathological stage in APP^NL-F/NL-F^ mice. At this age, both aggregated and soluble Aβ42 levels increase sharply, coinciding with plaque deposition, the onset of gliosis, and impairment of spatial memory. These findings refine the temporal framework of the model and reveal a coordinated progression from molecular alterations to cellular activation and early functional decline.

A key finding of this study is the close temporal alignment between this early molecular–functional transition and neuroinflammatory changes, which are central to AD pathogenesis ^38^. Here, we show that the onset of microgliosis and astrogliosis is accompanied by increased expression of genes associated with the disease-associated microglia (DAM) state, including *Cst7*, *Clec7a*, *Tyrobp* and *Trem2* ^8^, which are implicated in amyloid handling and debris clearance ^39–41^. By 12 months, when amyloidosis is established, this DAM signature is further amplified.

Single-cell profiling of microglia from 12-month-old APP^NL-F/NL-F^ mice and controls revealed eight distinct microglial states. In homozygous mice, this landscape was significantly remodeled, with an expansion of senescent-like (C2) and lipid-metabolic microglia (C9) states at the expense of homeostatic (C1 and C3) microglia, supporting an early convergence between aging-associated programs and amyloid-driven microglial reprogramming. Notably, amyloid-associated transcriptional changes were also detected within C1/C3 homeostatic clusters, indicating that early molecular perturbations precede overt phenotypic transitions.

Together, these observations indicate that the APP^NL-F/NL-F^ model captures not only amyloid-driven gliosis, but also mirrors age-like trajectories of microglial dysfunction, providing a relevant framework for studying the interaction between aging and AD-related pathology, a processes that is currently being evaluated in humans ^42^. This is particularly relevant given growing evidence that microglial senescence contributes to AD pathogenesis ^43^. Aging-related vulnerability of senescent-like microglia to amyloidosis may provide a rationale for exploring senolytic or senomorphic therapeutic strategies ^44^.

Our in depth study also brings forward the cognitive spatial memory impairments in APP^NL-F/NL-F^ mice to 9 months, from the previously reported deficits at 12 to 18 months ^12,19,37,45,46^. We were likely able to detect earlier phenotypes due to our use of low-stress, non-aversive behavioural paradigms. Indeed, other studies have also hinted at earlier behavioural alterations, including compulsivity ^19^, changes in functional connectivity ^21^, and motivational deficits ^20^.

Women are more vulnerable to AD ^47,48^ and sexual dimorphism is well established in microglial biology ^49^. However, sex-specific effects remain largely underexplored in the APP^NL-F^ model, especially at the early stages of the disease. While we observed similar amyloid burden, gliosis, and behavioural outcomes in males and females in the APP^NL-F^ model, our transcriptional analyses revealed a more pronounced microglial response in females. Yet, the functional implications of these transcriptional differences remain to be established.

Finally, while heterozygous APP^NL-F/WT^ mice lack overt plaque and glial pathology, the presence of early spatial memory impairments in these animals suggests that elevated soluble Aβ species may be sufficient to induce mild spatial memory impairment. Thus, the homozygous APP^NL–F/NL–F^ model provides a robust platform for studying plaque-associated pathology, whereas the heterozygous model may be informative for investigating plaque-independent, soluble Aβ–mediated toxicity.

In conclusion, our study refines the temporal framework of early pathology in APP^NL-F/NL-F^ mice, and identifies a critical inflection point during midlife at which amyloid deposition, microglial state transitions, and early cognitive decline converge. By aligning microglial state transitions and early neuroinflammation with amyloid escalation and emerging spatial memory decline, this work highlights a mechanistic axis through which aging and amyloidosis intersect and points to an early therapeutic window in which targeting microglial dysfunction may have maximal impact.

## Limitations

Although the single-cell data generated in this work enables the exploration of microglial diversity, its statistical power was limited by the use of a single library per genotype. Nonetheless, transcriptomic findings were consistent with qPCR data, supporting their robustness. Further, because our single-cell transcriptomic analysis was performed at 12 months, i.e. three months after the initial pathological inflection point, the observed changes in microglial populations represent an established disease state. This study cannot definitively conclude whether these microglial shifts drive the onset of pathology at 9 months or emerge as a consequence of it; this will be the focus of future studies. Finally, environmental and facility-specific factors, the experimental “exposome”, may influence the precise timing of pathology. While we consistently observed an inflection point during midlife, the exact age at which it occurs may vary across breeding conditions. Importantly, however, the convergence of molecular, cellular, and behavioural alterations across independent experimental readouts strengthens the robustness of our conclusions and the translational relevance of the APP^NL–F/NL–F^ model of midlife changes for presymptomatic inflection points in human middle age.

## Supplementary table legends

**Supplementary Table 1:**
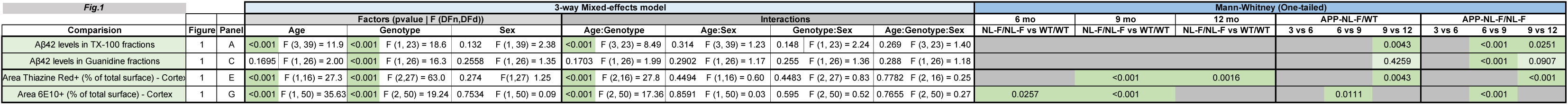
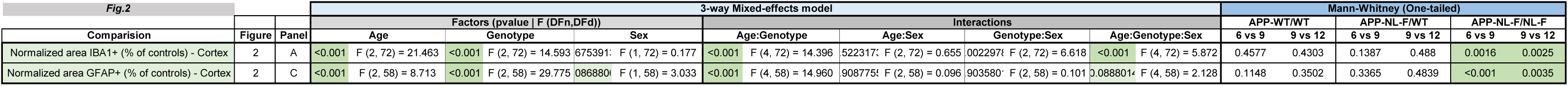
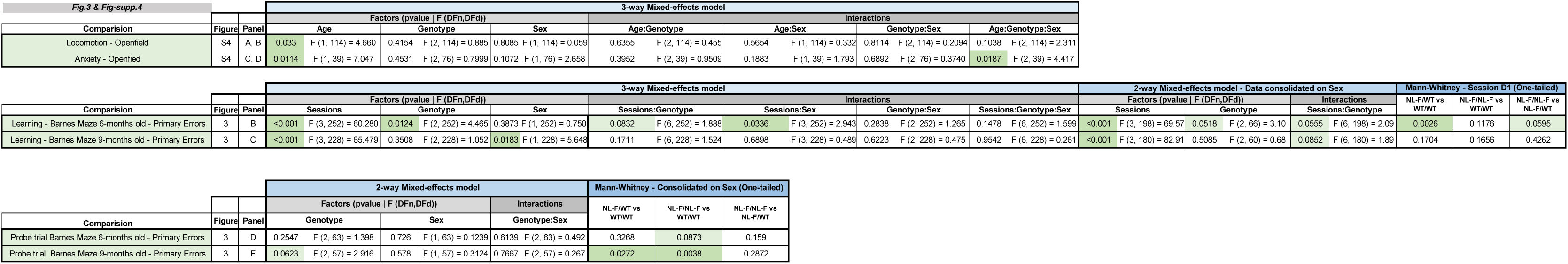
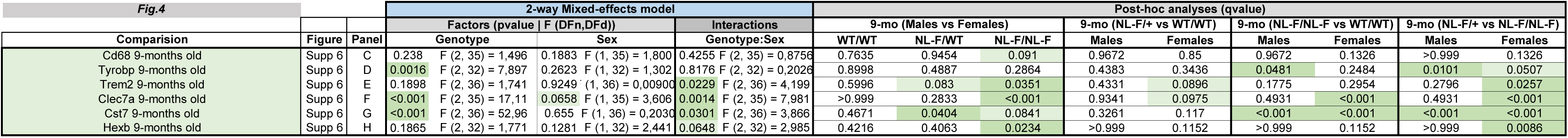
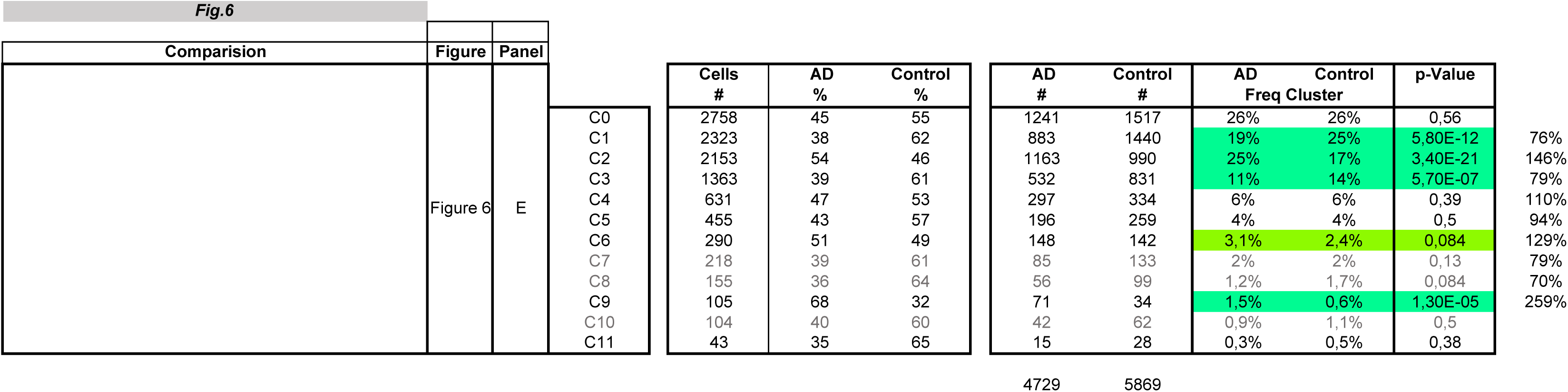
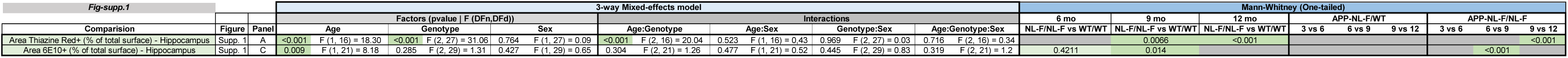
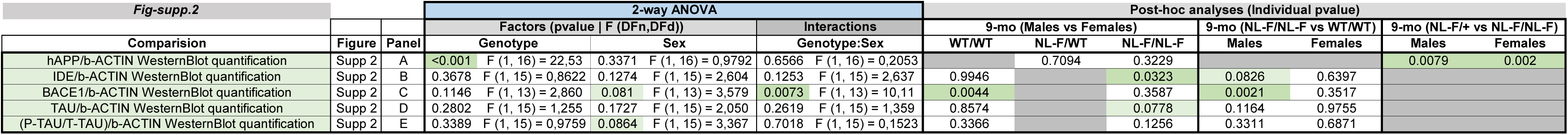
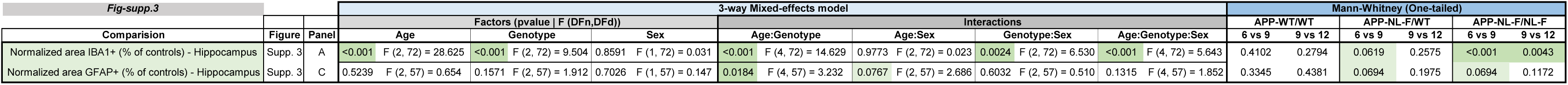
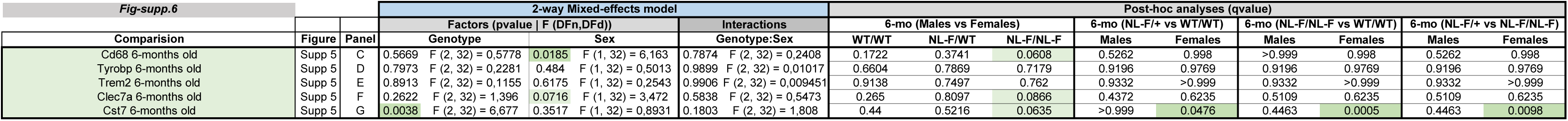
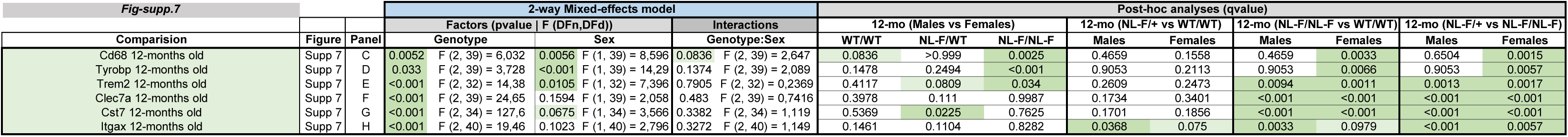
Detailed results of all statistical analyses performed in this study. Each sheet corresponds to a specific main or supplementary figure.

**Supplementary Table 2.**
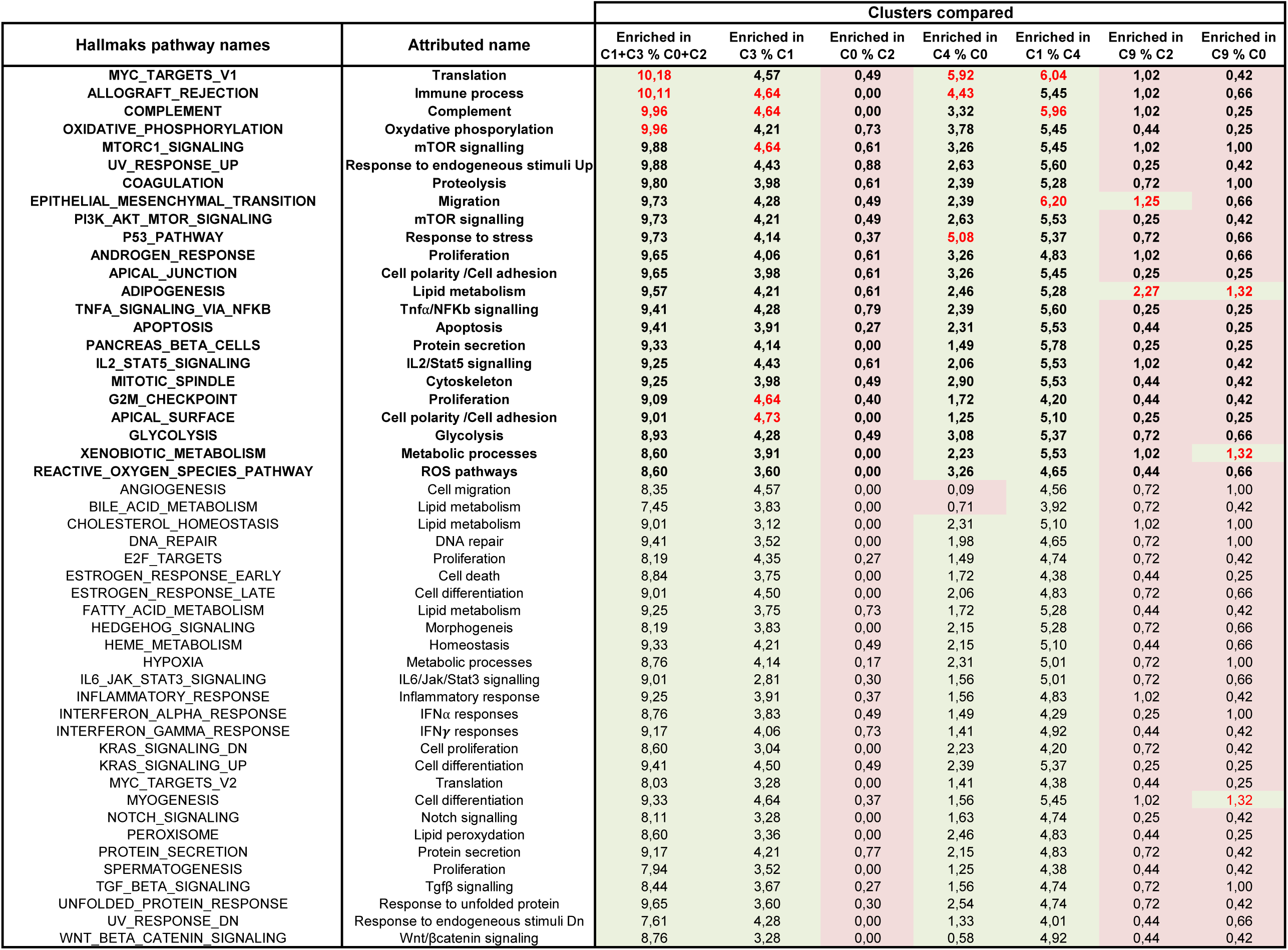
(related to Fig. 6D): Single-cell pathway analysis (SCPA), selected cluster comparisons in APP^WT/WT^ cells (Q values). Green cells: significant deregulation (Q<0.05); Red: non-significant deregulation. Bold: Union of Top 10 pathways of each comparison. Red labels: top three up-regulated per comparison. Genes from each Hallmark pathway were retrieved and analysed by GO enrichment to assign pathway names that better reflect microglia-specific biological processes.

**Supplementary Table 3.** (related to Fig. 6F): single-cell pathway analysis (SCPA) between APP^WT/WT^ and APP^NL-F/NL^ ^F^ within each cluster (Q values). Pathway renaming was performed as in Supplementary Supp-Table 2.

| Hallmark pathways | Relevant related pathways | C0 | C1 | C2 | C3 | C4 | C5 | C6 | C9 |
| --- | --- | --- | --- | --- | --- | --- | --- | --- | --- |
| PANCREAS BETA CELLS | Protein secretion | 3,03 | 3,92 | 3,97 | 5,82 | 0,83 | 0,00 | 2,07 | 0,01 |
| UV RESPONSE DN | Response to endogenous stimuli DN | 8,36 | 7,28 | 8,44 | 8,95 | 3,95 | 0,00 | 1,95 | 1,84 |
| ESTROGEN RESPONSE EARLY | Cell death | 8,12 | 8,09 | 8,51 | 8,95 | 4,09 | 0,00 | 1,65 | 1,84 |
| KRAS SIGNALING DN | Cell proliferation | 8,51 | 7,77 | 8,28 | 9,43 | 4,81 | 3,42 | 2,17 | 2,33 |
| NOTCH SIGNALING | Notch signalling | 6,40 | 5,79 | 6,47 | 7,63 | 2,60 | 1,21 | 1,34 | 1,03 |
| APICAL SURFACE | Cell polarity /Cell adhesion | 6,33 | 4,82 | 6,55 | 5,41 | 2,64 | 1,73 | 0,82 | 1,03 |
| WNT BETA CATENIN SIGNALING | Wnt/bcatenin signaling | 5,11 | 3,51 | 5,02 | 4,01 | 1,93 | 1,11 | 0,45 | 1,12 |
| ANGIOGENESIS | Cell Migration | 3,42 | 3,36 | 2,54 | 2,78 | 0,33 | 0,00 | 0,00 | 1,40 |
| HEDGEHOG SIGNALING | Morphogenesis | 6,25 | 4,57 | 5,41 | 6,15 | 2,29 | 0,00 | 0,45 | 1,84 |
| ANDROGEN RESPONSE | Proliferation | 6,65 | 5,39 | 6,71 | 6,47 | 1,34 | 0,00 | 0,00 | 1,03 |
| MYOGENESIS | Cell differentiation | 6,65 | 5,39 | 6,15 | 6,22 | 0,79 | 0,00 | 0,00 | 1,26 |
| ADIPOGENESIS | Lipid metabolism | 4,62 | 1,56 | 3,17 | 0,00 | 0,00 | 0,00 | 0,00 | 0,01 |
| INFLAMMATORY RESPONSE | Inflammatory response | 5,03 | 1,34 | 4,13 | 0,00 | 0,00 | 0,00 | 0,00 | 0,01 |
| G2M CHECKPOINT | Proliferation | 3,50 | 1,19 | 2,94 | 0,00 | 0,00 | 0,00 | 0,00 | 0,01 |
| HYPOXIA | Metabolic processes | 5,19 | 1,71 | 4,05 | 0,00 | 0,00 | 0,00 | 0,00 | 0,01 |
| EPITHELIAL MESENCHYMAL TRANSITION | Cell Migration | 4,38 | 1,19 | 3,65 | 0,55 | 0,00 | 0,00 | 0,00 | 0,01 |
| PI3K AKT MTOR SIGNALING | mTOR signalling | 4,62 | 1,71 | 4,21 | 0,62 | 0,00 | 0,00 | 0,00 | 0,01 |
| P53 PATHWAY | Response to stress | 4,30 | 2,48 | 3,02 | 0,00 | 0,00 | 0,00 | 0,00 | 0,01 |
| BILE ACID METABOLISM | Lipid metabolism | 3,90 | 1,48 | 3,57 | 0,00 | 0,47 | 0,00 | 0,00 | 0,01 |
| HEME METABOLISM | Homeostasis | 4,62 | 2,10 | 4,21 | 0,00 | 0,00 | 0,00 | 0,00 | 0,01 |
| IL6 JAK STAT3 SIGNALING | IL6/Jak/Stat3 signalling | 3,11 | 1,34 | 3,09 | 0,00 | 0,00 | 0,00 | 0,00 | 0,01 |
| ESTROGEN RESPONSE LATE | Cell differentiation | 6,01 | 2,40 | 5,25 | 2,62 | 0,00 | 0,00 | 0,00 | 0,01 |
| MITOTIC SPINDLE | Cytoskeleton | 6,72 | 3,43 | 5,74 | 2,54 | 0,00 | 0,00 | 0,00 | 0,01 |
| IL2 STAT5 SIGNALING | IL2/Stat5 signalling | 5,27 | 2,55 | 5,17 | 2,47 | 0,00 | 0,00 | 0,00 | 0,01 |
| TGF BETA SIGNALING | Tgfb signalling | 5,43 | 2,48 | 4,77 | 2,78 | 0,00 | 0,00 | 0,00 | 0,29 |
| KRAS SIGNALING UP | Cell differentiation | 6,09 | 2,10 | 4,36 | 1,68 | 0,00 | 0,00 | 0,00 | 0,71 |
| APICAL JUNCTION | Cell polarity /Cell adhesion | 6,09 | 2,55 | 4,28 | 1,68 | 0,00 | 0,00 | 0,00 | 0,01 |
| COMPLEMENT | Complement | 6,33 | 2,17 | 5,02 | 1,82 | 0,00 | 0,00 | 0,00 | 0,01 |
| SPERMATOGENESIS | Proliferation | 4,06 | 2,10 | 4,21 | 0,79 | 0,00 | 0,00 | 0,00 | 0,01 |
| ALLOGRAFT REJECTION | Regulation of Immune system process | 5,43 | 2,32 | 5,02 | 1,68 | 0,00 | 0,00 | 0,00 | 0,01 |
| UV RESPONSE UP | Response to endogenous stimuli Up | 6,79 | 3,03 | 5,58 | 1,90 | 0,00 | 0,00 | 0,00 | 0,01 |
| MYC TARGETS V1 | Translation | 1,64 | -0,31 | 0,04 | 0,00 | 0,00 | 0,00 | 0,00 | 0,01 |
| MTORC1 SIGNALING | mTORC1 signalling | 2,09 | 0,00 | 0,23 | 0,00 | 0,00 | 0,00 | 0,00 | 0,01 |
| OXIDATIVE PHOSPHORYLATION | Oxydative phosphorylation | 0,93 | 0,00 | 0,04 | 0,00 | 0,00 | 0,00 | 0,00 | 0,01 |
| COAGULATION | Proteolysis | 3,11 | 0,00 | 0,97 | 0,00 | 0,00 | 0,00 | 0,00 | 0,01 |
| FATTY ACID METABOLISM | Lipid metabolism | 4,22 | 0,38 | 1,55 | 0,00 | 0,00 | 0,00 | 0,00 | 0,01 |
| E2F TARGETS | Proliferation | 1,94 | 0,00 | 0,91 | 0,00 | 0,00 | 0,00 | 0,00 | 0,01 |
| INTERFERON ALPHA RESPONSE | Ifna response | 3,58 | 0,00 | 1,55 | 0,00 | 0,00 | 0,00 | 0,00 | 0,01 |
| REACTIVE OXYGEN SPECIES PATHWAY | ROS pathway | 2,87 | 0,00 | 3,65 | 0,00 | 0,00 | 0,00 | 0,00 | 0,01 |
| GLYCOLYSIS | Glycolysis | 4,22 | 0,67 | 3,49 | 0,00 | 0,00 | 0,00 | 0,00 | 0,01 |
| PROTEIN SECRETION | Protein secretion | 3,82 | 0,60 | 3,33 | 0,00 | 0,00 | 0,00 | 0,00 | 0,01 |
| PEROXISOME | Lipid peroxydation | 3,98 | 0,24 | 3,02 | 0,00 | 0,00 | 0,00 | 0,00 | 0,01 |
| UNFOLDED PROTEIN RESPONSE | Response to unfolded protein | 2,09 | 0,16 | 1,70 | 0,00 | 0,00 | 0,00 | 0,00 | 0,01 |
| TNFA SIGNALING VIA NFKB | Tnfa/NFkb signalling | 2,40 | 0,00 | 1,70 | 0,00 | 0,00 | 0,00 | 0,00 | 0,01 |
| DNA REPAIR | DNA repair | 0,99 | 0,00 | 0,78 | 0,00 | 0,00 | 0,00 | 0,00 | 0,01 |
| XENOBIOTIC METABOLISM | Metabolic processes | 2,55 | 0,00 | 1,92 | 0,00 | 0,00 | 0,00 | 0,00 | 0,01 |
| CHOLESTEROL HOMEOSTASIS | Lipid metabolism | 3,58 | 0,00 | 3,41 | 0,00 | 0,00 | 0,00 | 0,00 | 0,01 |
| INTERFERON GAMMA RESPONSE | Ifng response | 3,34 | 0,00 | 2,86 | 0,00 | 0,00 | 0,00 | 0,00 | 0,01 |
| APOPTOSIS | Apoptosis | 2,48 | 0,00 | 1,70 | 0,00 | 0,00 | 0,00 | 0,00 | 0,01 |
| MYC TARGETS V2 | Translation | 2,48 | 0,00 | 2,07 | 0,00 | 0,00 | 0,00 | 0,00 | 0,01 |

## Acknowledgements

The authors thank iExplore, Montpellier Génomique (MGX) and Montpellier Rio Imaging (MRI) plateforms for their assistance in data acquisition and analysis. We also thank Drs V. Compan, E. Audinat, M. Weimershaus and J.P. Venables (Science Sense) for their critical review during manuscript preparation. Some of the figures were created with https://BioRender.com under the academic user license of HH.

## Author contributions

MP: Data collection and analysis, Drafting the figures; Writing original draft. ALHG: Data collection and analysis, Writing original draft; VB: scRNA-seq analysis; Drafting figures; VG: scRNA-seq experiments; PT: qPCR experiments; NL: Mouse Genotyping, Ressources management; ML: Behavioral testing; AV: Western-Blot experiments and analysis; DS: Technical support for single cell technology, Library construction; VP: Western-blot supervision, Manuscript revision; FAR: Funding acquisition & discussion; MD: Result analysis & discussion, Manuscript revision; ST: APP^NL-F^ generous gift, Manuscript revision; ST: APP^NL-F^ generous gift, Manuscript revision; HH: Conceptualization, Supervision; Resources, Funding acquisition, Project administration, Writing original draft.

## Data Availability

The scRNA-seq dataset analyzed in this study will be submitted to the GEO repository upon acceptance of the manuscript. Other data supporting the findings are available from the corresponding author upon reasonable request.

## Supplementary figure legends

**Supplementary Figure 1:**
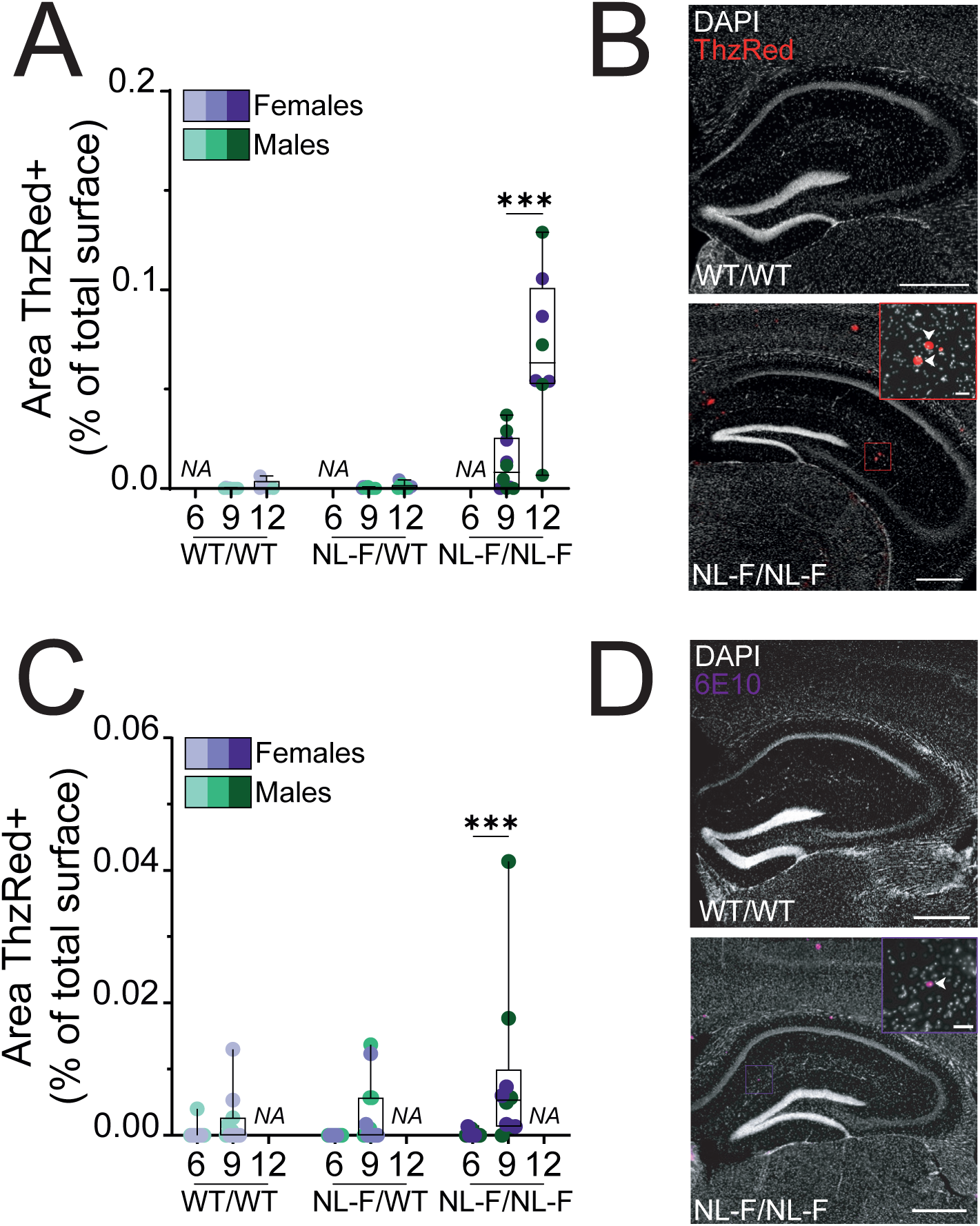
Aβ accumulate early in a genotype-dependant manner in the hippocampus of APP^WT/NL-F^ and APP^NL-F/NL-F^ mice – related to Figure 1. (**A, C**) Quantification of Thiazine Red (ThzR)-positive (**A**) and 6E10-positive areas (**C**) in the hippocampus of APP^NL-F^ mice aged 3, 6, 9 and 12 months. (**B, D**) Representative images of ThzR staining in 12-mo mice (**B, red**) and 9-mo mice (**D, purple**) from APP^WT/WT^ and APP^NL-F/NL-F^ genotypes. Arrow heads indicate amyloid deposits. Scale bar: 500 µm (insert: 50µm). Data are presented as box-and-whisker plots showing the median, interquartile range (25^th^–75^th^ percentiles), and minimum and maximum values. Each data point represents one animal. Sex is indicated by color (males, green; females, purple), while genotype is indicated by color shade (APP^WT/WT^, light; APP^NL-F/WT^, medium; APP^NL-F/NL-F^, dark). <u>Statistical analyses on the figure</u>: Mann-Whitney one-tailed between consecutive age groups within the same genotype. *** p<0.001. “NA”: Not Analyzed.

**Supplementary Figure 2:**
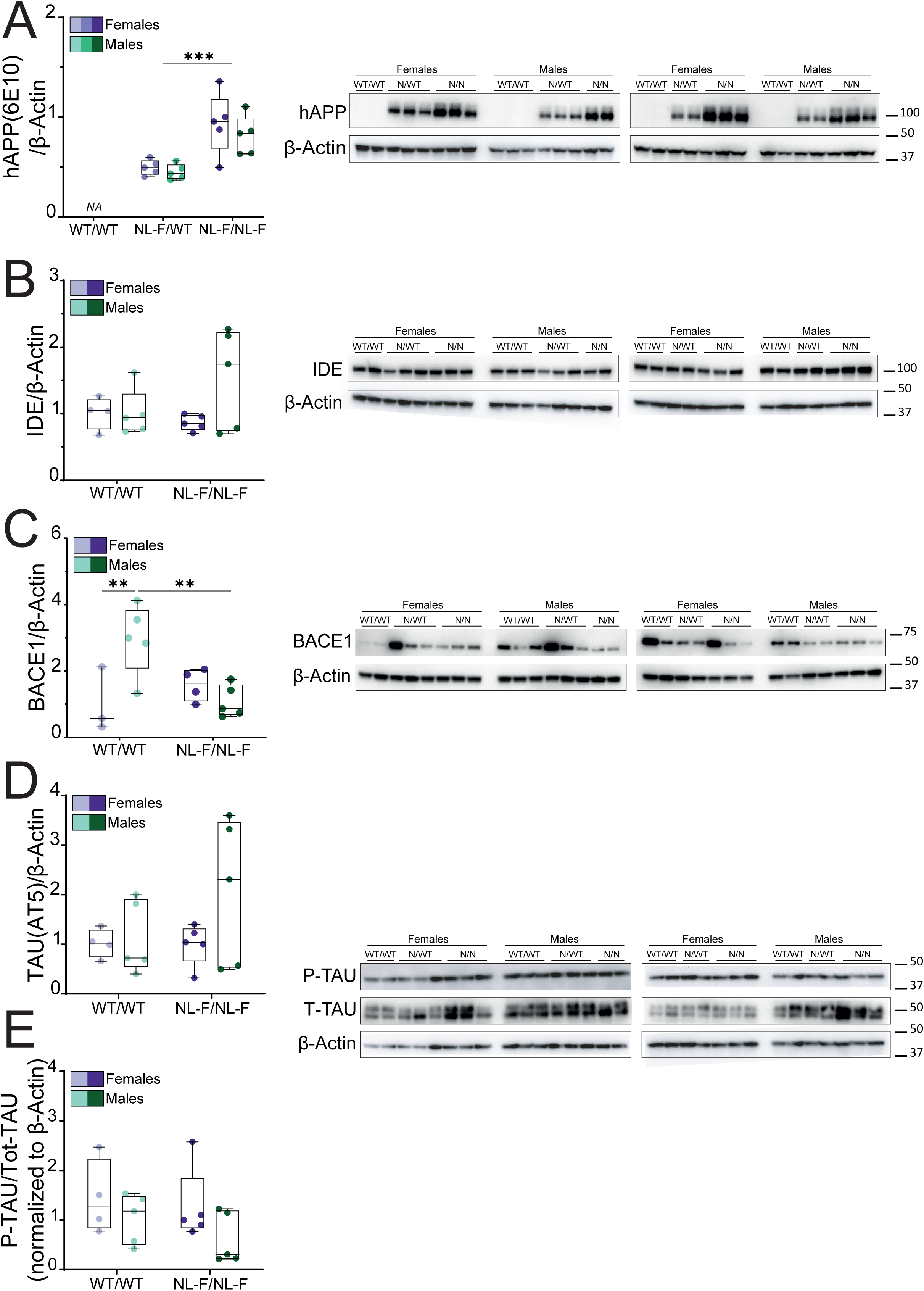
Western blot quantification of Alzheimer’s disease-related proteins in cortical extracts of 9-mo APP^NL-F^ mice. (**A**) Quantification of hAPP (6E10) protein levels in cortical extracts from APP^WT/WT^, APP^NL-F/WT^ and APP^NL-F/NL-F^ mice. (**B-D**) Quantification of insulin-degrading enzyme (IDE, **B**), β-secretase 1 (BACE1, **C**) and TAU (**D**) protein levels in cortical extracts from APP^WT/WT^ and APP^NL-F/NL-F^ mice. (**E**) Quantification of phosphorylated TAU (P-TAU, AT8) normalized to total TAU (AT5) levels in cortical extracts from APP^WT/WT^ and APP^NL-F/NL-F^ mice. (**A-E**) Densitometric values were normalized to β-actin (A–D) or total TAU (E) and are presented as box-and-whisker plots showing the median, interquartile range (25^th^–75^th^ percentiles), and minimum and maximum values. Each data point represents one animal. Sex is indicated by color (males, green; females, purple), while genotype is indicated by color shade (APP^WT/WT^, light; APP^NL-F/WT^, medium; APP^NL-F/NL-F^, dark). Corresponding Western blot membranes are shown. <u>Statistical analysis</u>: Two-way ANOVA with genotype and sex as between-subject factors; FDR-corrected post-hoc test: ** p< 0.01, *** p<0.001.

**Supplementary Figure 3:**
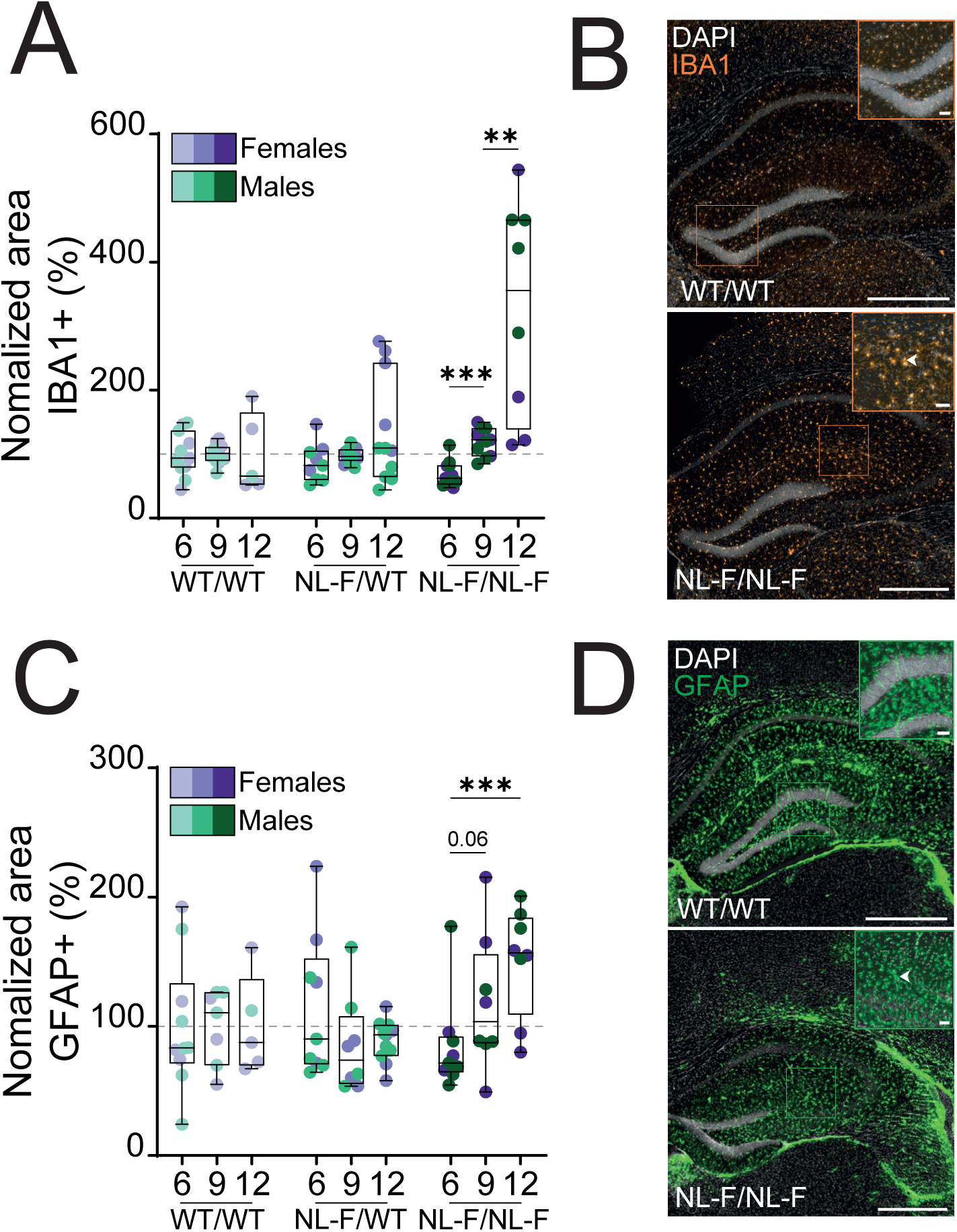
Progressive increase in microgliosis and astrogliosis in the hippocampus of APP^NL-F/NL-F^ mice during the development of amyloidosis. Related to Figure 2. (**A**) Quantification of hippocampal-IBA1 (microglia) positive area in 6-, 9- and 12-month-old APP^NL-F^ mice, normalized to aged matched controls. (**B**) Representative IBA1 immunostaining of 12-mo APP^WT/WT^ and APP^NL-F/NL-F^ mice. Arrow head in insert shows microglia clustering. Scale bar: 500µm; insert: 50µm. (**C**) Quantification of hippocampal-GFAP positive area in 6-, 9- and 12-month-old APP^NL-F^ mice, normalized to aged matched controls. (**D**) Representative GFAP immunostaining of 12-mo APP^WT/WT^ and APP^NL-F/NL-F^ mice. Arrow head in insert shows astrocyte clustering. Scale bar: 500µm; insert: 50µm. Quantitative data are shown as box-and-whisker plots indicating the median, interquartile range (25th–75th percentiles), and minimum and maximum values. Each data point represents one animal. Sex is indicated by color (males, green; females, purple), while genotype is indicated by color shade (APP^WT/WT^, light; APP^NL-F/WT^, medium; APP^NL-F/NL-F^, dark). Horizontal dotted line represents 100% of the mean control value. <u>Statistical analyses on the figures</u>: one-tailed Mann-Whitney between ages within a same genotype. ** p<0.01, *** p<0.001.

**Supplementary Figure 4:**
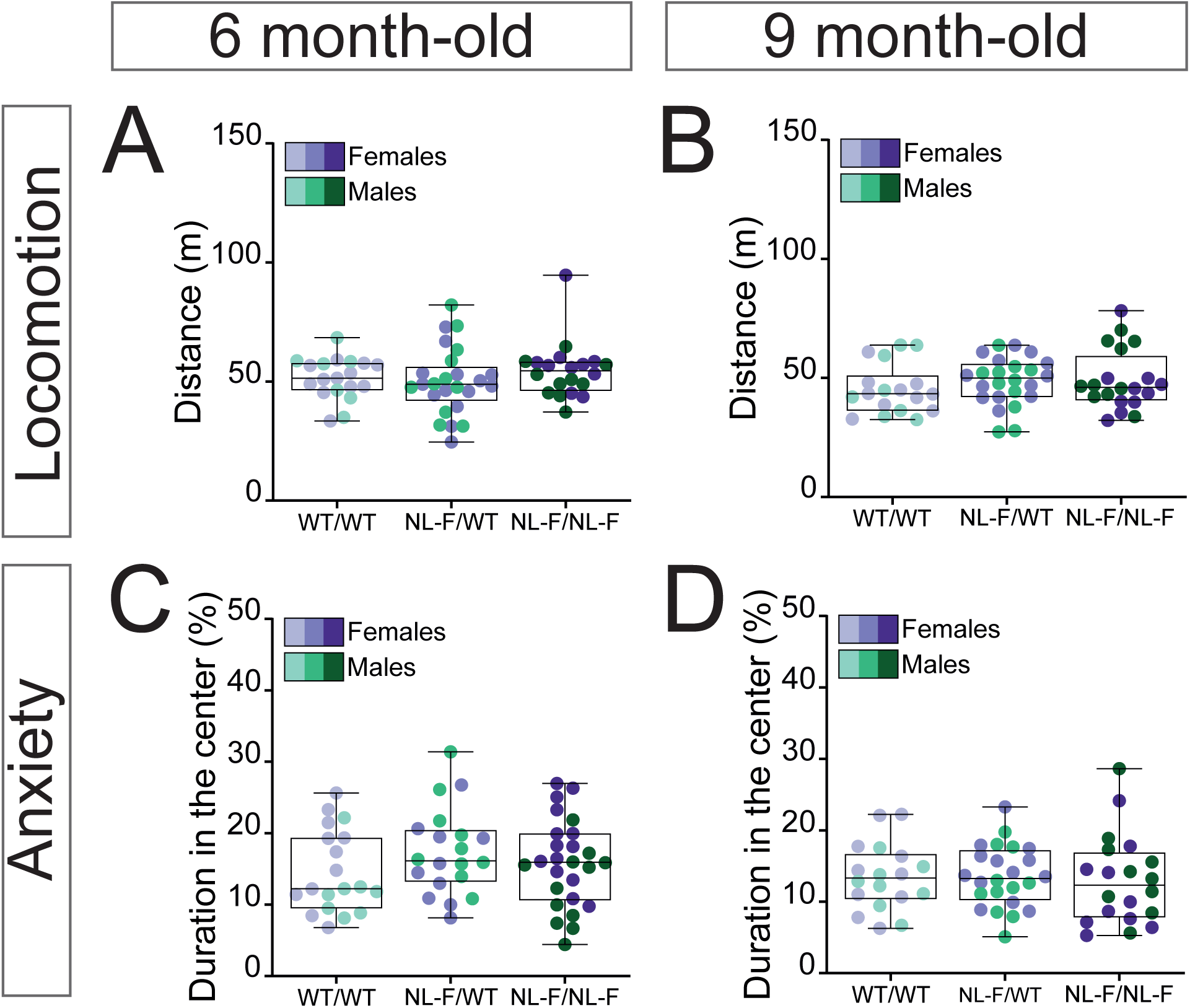
Complementary behavioural analyses in 6- and 9-mo mice. Related to Figure 3. (**A**, **B**) Locomotor activity of 6-mo (**A**) and 9-mo (**B**) mice in the open field test. (**C**, **D**) Anxiety-like behavior of 6-mo (**C**) and 9-mo (**D**) APP^NL-F^ mice in the open field test, are represented as box-and-whisker plots indicating the median, interquartile range (25th–75th percentiles), and minimum and maximum values. Each data point represents one animal. Sex is indicated by color (males, green; females, purple), while genotype is indicated by color shade (APP^WT/WT^, light; APP^NL-F/WT^, medium; APP^NL-F/NL-F^, dark).

**Supplementary Figure 5:**
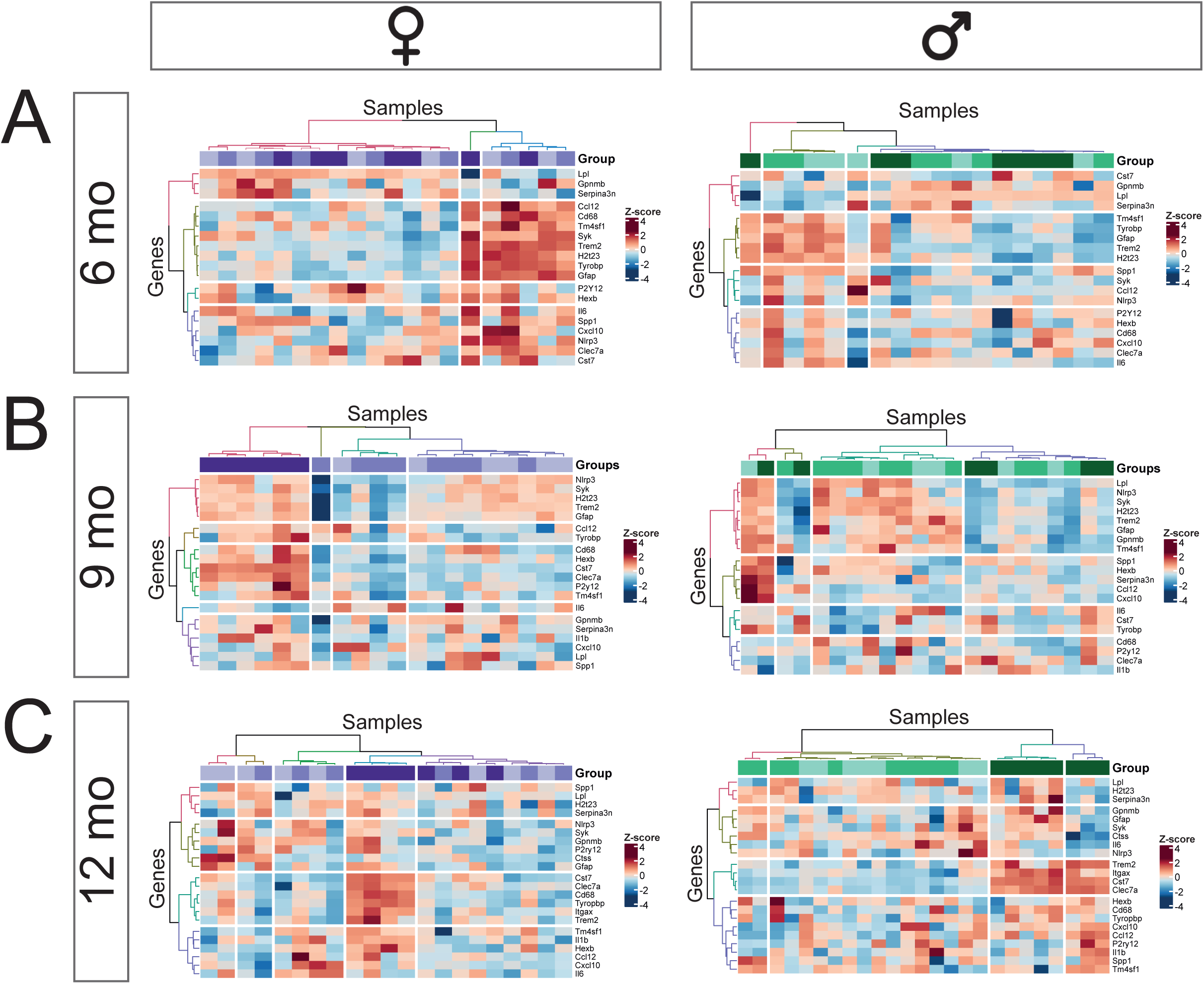
Gene expression changes in the 6-mo mice cortical homogenates for a range of inflammation and AD-related gens. Related to Figure 4. (**A-C**) Heatmaps of differentially expressed genes in APP^NL-F^ mice across pathology progression (6, 9 and 12-mo). Data are presented separately for males (Right column) and females (left column). Sex is indicated by color (males, green; females, purple), while genotype is indicated by color shade (APP^WT/WT^, light; APP^NL-F/WT^, medium; APP^NL-F/NL-F^, dark). Scaled expression values (row Z scores) are shown using a blue–red color scale, with red indicating higher and blue lower expression.

**Supplementary Figure 6:**
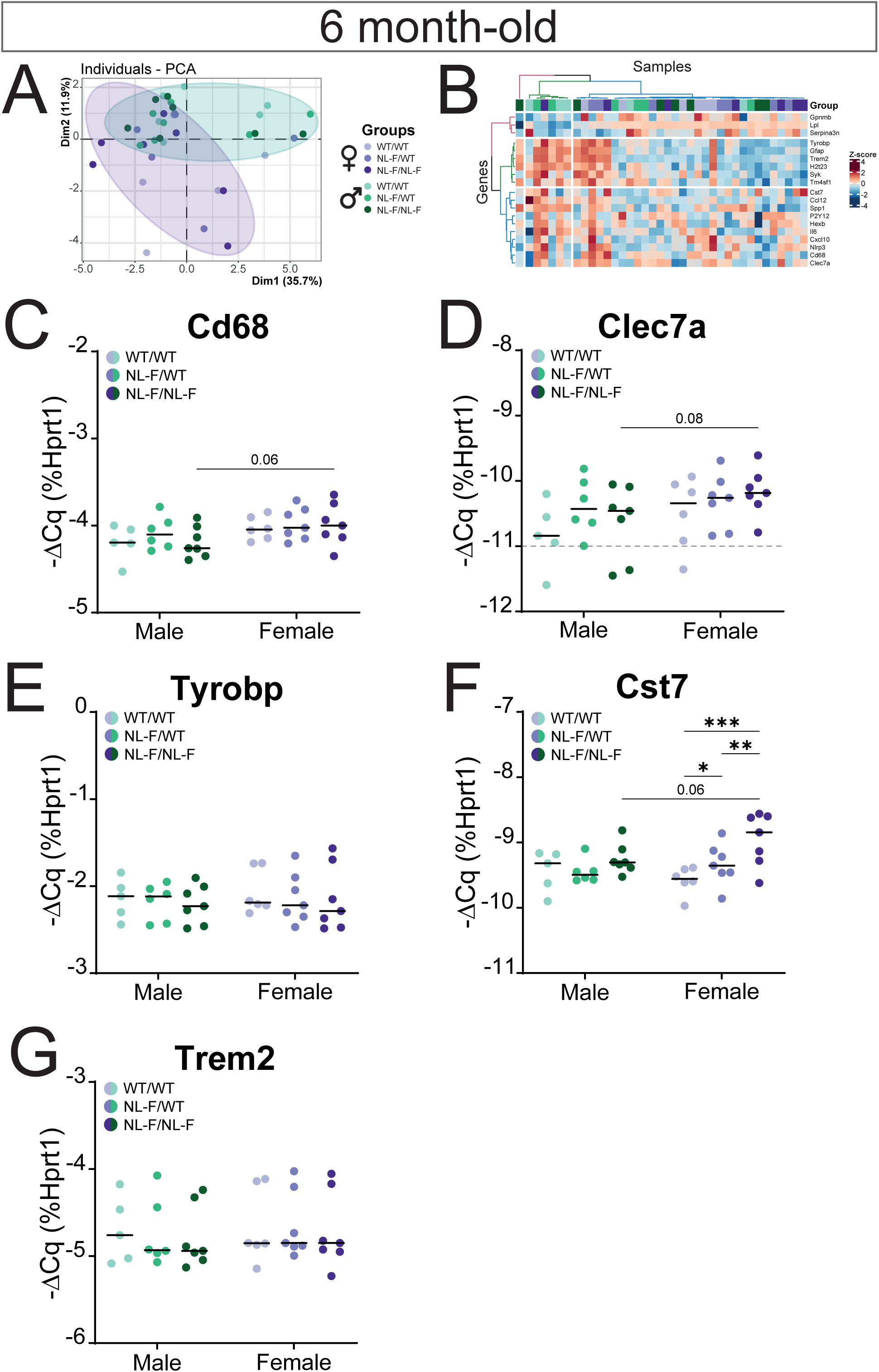
Gene expression changes in the 6-mo mice cortical homogenates for a range of inflammation and AD-related gens. Related to Figure 4. (**A**) PCA plot of individual samples projected along PC1 and PC2. (**B**) Heatmap of differentially expressed genes. Scaled expression values (row Z score) are shown using a blue–red color scale with red indicating higher expression, and blue lower expression. (**C**-**G**) Gene expression changes measured by qPCR and plotted as −ΔCq for a selection of genes. Data are shown as dot plots indicating the median. Each data point represents one animal. Sex is indicated by color (males, green; females, purple), while genotype is indicated by color shade (APP^WT/WT^, light; APP^NL-F/WT^, medium; APP^NL-F/NL-F^, dark). <u>Statistical analysis</u> for each selected gene: 2-way ANOVA with genotype and sex as between-subject factors. FDR corrected post-hoc tests. * p<0.05.

**Supplementary Figure 7:**
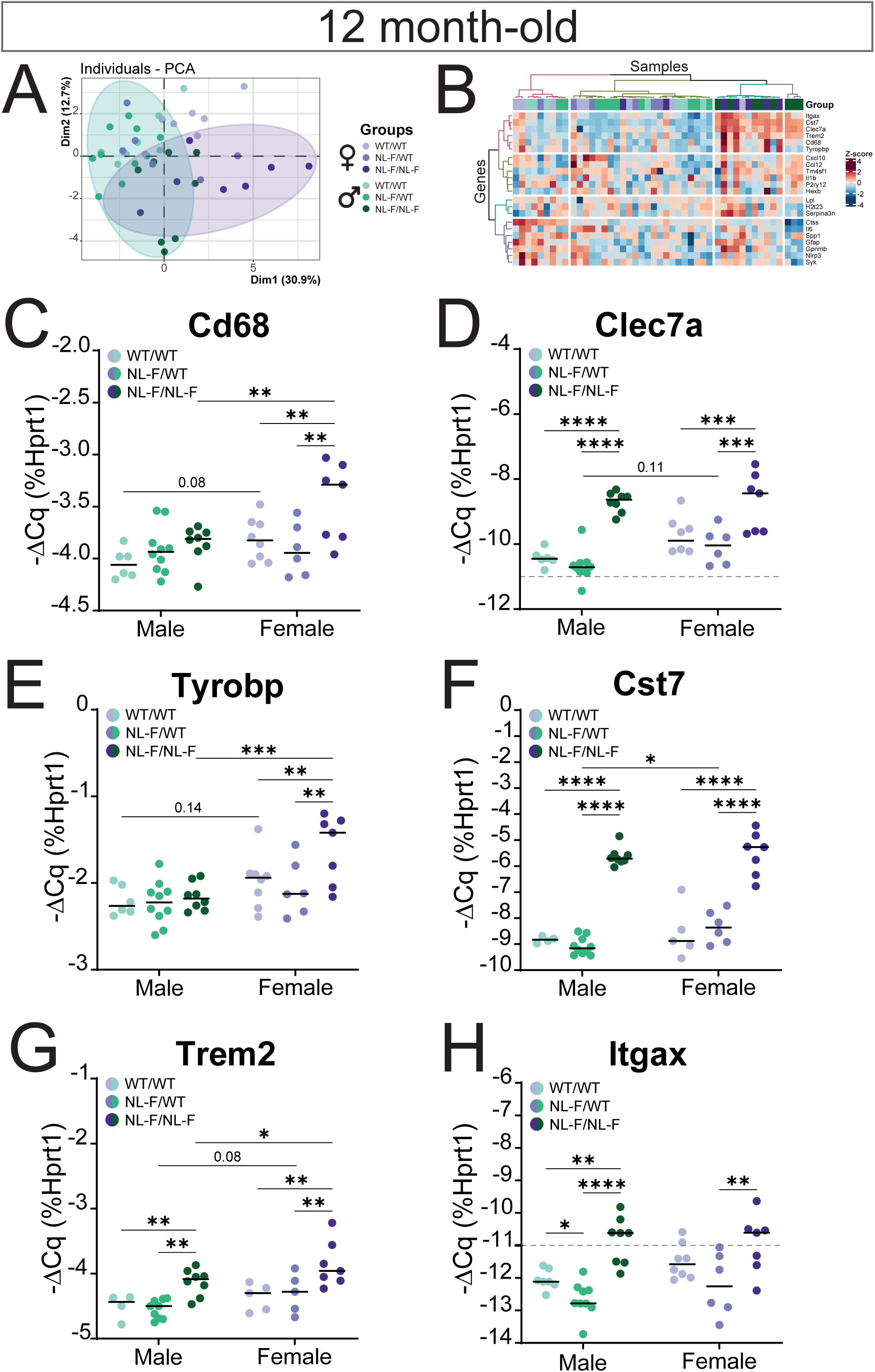
Gene expression changes in the 12-mo mice cortical homogenates for a range of inflammation and AD-related gens. Related to Figure 5. (**A**) PCA plot of individual samples projected along PC1 and PC2. (**B**) Heatmaps of differentially expressed genes in APP^WT/WT^, APP^NL-F/WT^ and APP^NL-F/NL-F^ mice. Scaled expression values (row Z score) are shown using a blue–red color scale with red indicating higher expression, and blue lower expression. (**C**-**G**) Gene expression changes measured by qPCR and plotted as −ΔCq for a selection of genes. Data are shown as dot plots indicating the median. Each data point represents one animal. Sex is indicated by color (males, green; females, purple), while genotype is indicated by color shade (APP^WT/WT^, light; APP^NL-F/WT^, medium; APP^NL-F/NL-F^, dark). <u>Statistical analysis</u> for each selected gene: 2-way ANOVA with genotype and sex as between subjects’ factors. FDR corrected post-hoc tests. * p<0.05.

**Supplementary Figure 8:**
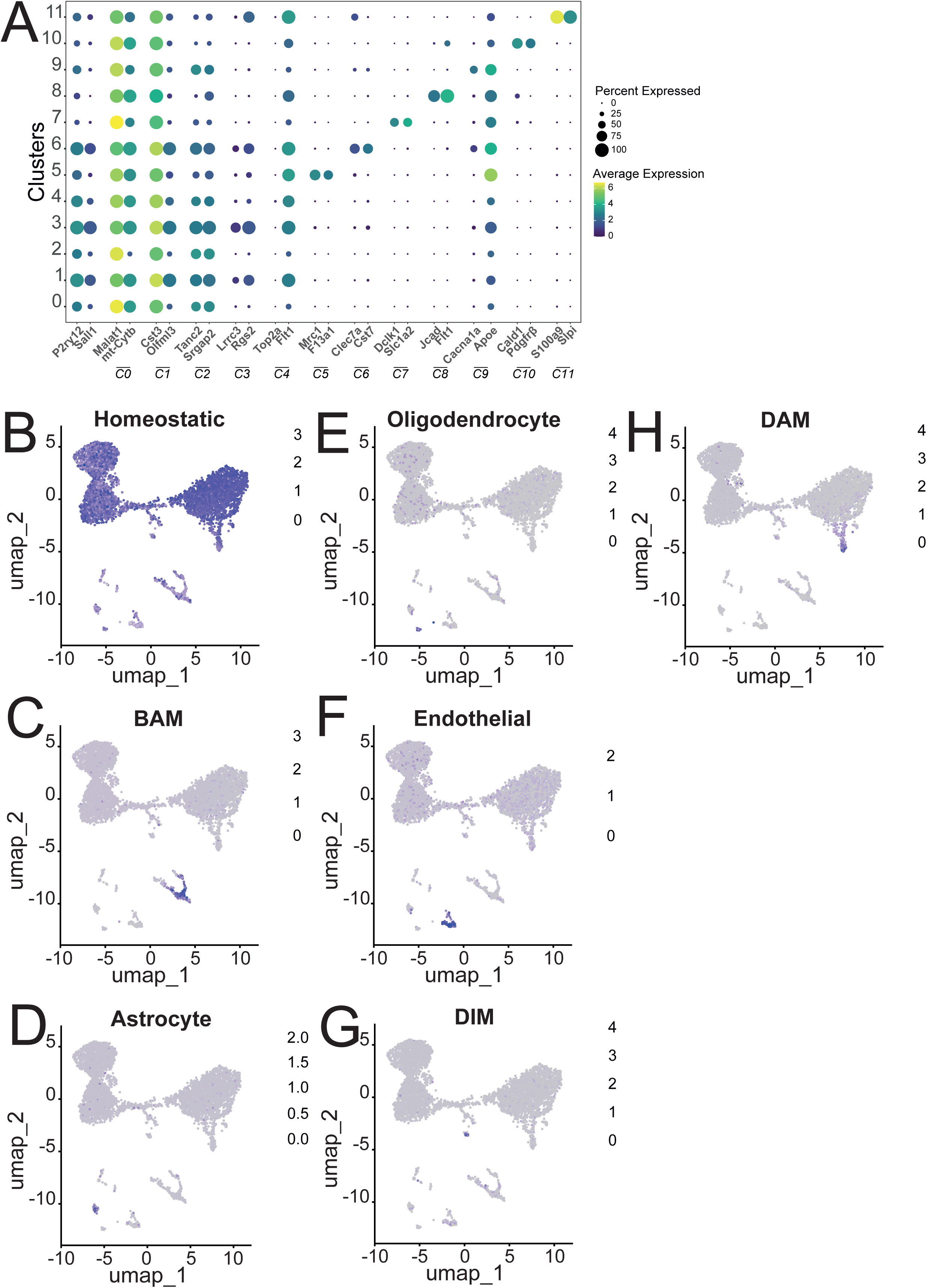
Characterisation of the different clusters. Related to Figure 6. (**A**) Dotplots showing representative marker gene expression for each cluster (**B-H**) UMAPs for specific molecular signatures in APP^WT/WT^ cels cells. (**A**) Homeostatic microglia (*Cx3cr1*, *Hexb*, *P2ry12*, *Sall1*, *Tmem119*); (**C**) Border Associated Macrophages (*Mrc1*, *Cd163*, *Lyve1*, *Ms4a7*); (**D**) Astrocytes (*Aqp4*, *Fgfr3*, *Gfap*, *Gja1*); (**E**) Oligodendrocytes (*Mobp*, *Mog*, *plp*1); (**F**) Endothelial cells (*Cldn5*, *Cdh5*, *Pecam1*, *Flt1*, *Vwf*); (**G**) Disease Inflammatory Macrophages (DIM); (**G**) Disease Associated Microglia (DAM).

**Supplementary Figure 9:**
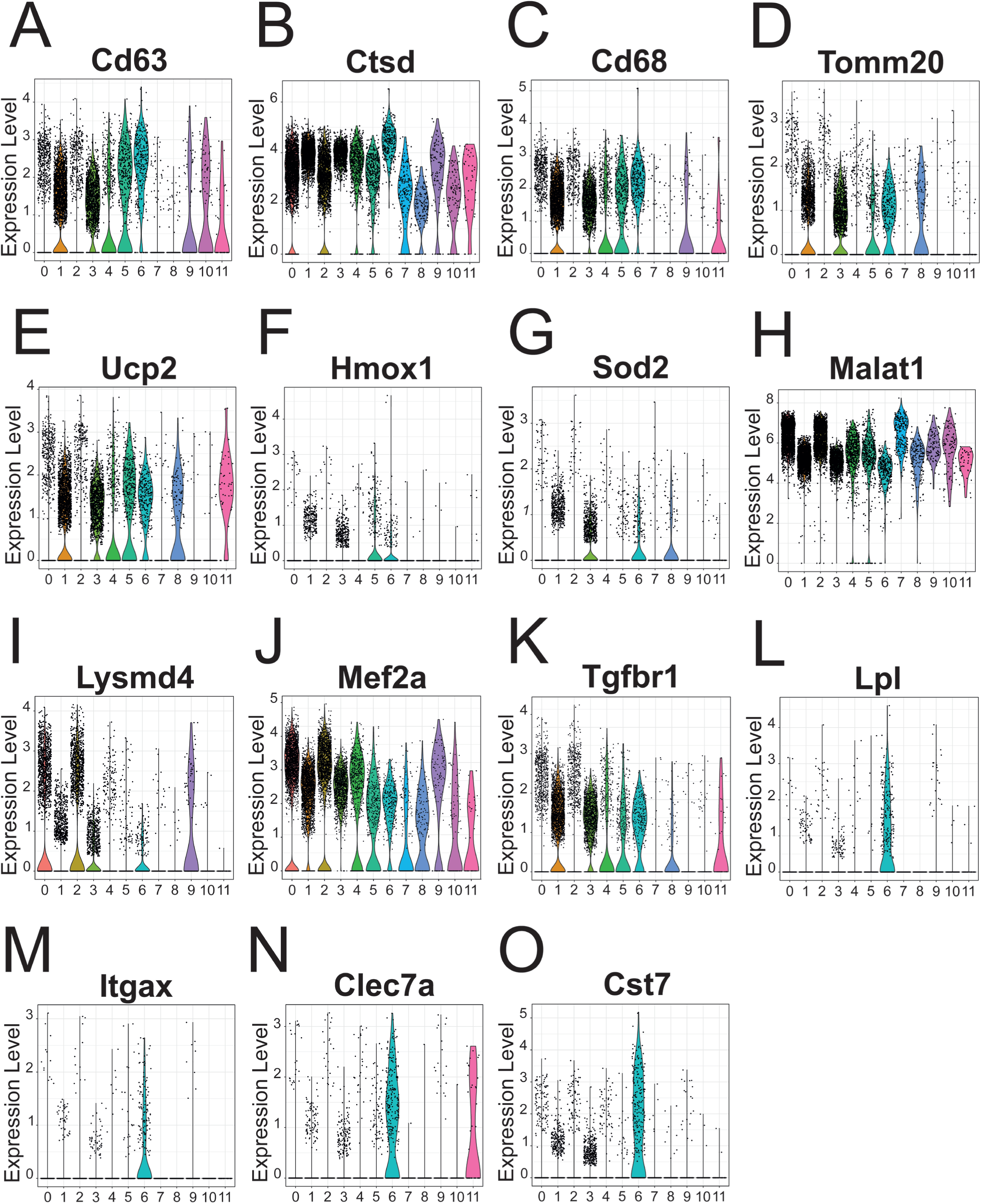
Characterisation of the different clusters. Related to Figure 6. Violin plots showing the normalized expression levels of cluster-defining genes described in the main text across the twelve microglial clusters identified by single-cell RNA sequencing.

## References

1. Heneka MT, Morgan D, Jessen F. Passive anti-amyloid β immunotherapy in Alzheimer’s disease—opportunities and challenges. The Lancet. 2024;404(10468):2198–2208. doi:10.1016/S0140-6736(24)01883-X

2. Braak H, Alafuzoff I, Arzberger T, Kretzschmar H, Del Tredici K. Staging of Alzheimer disease-associated neurofibrillary pathology using paraffin sections and immunocytochemistry. Acta Neuropathol (Berl). 2006;112(4):389–404. doi:10.1007/s00401-006-0127-z

3. Heneka MT, Carson MJ, El Khoury J, et al. Neuroinflammation in Alzheimer’s Disease. Lancet Neurol. 2015;14(4):388–405. doi:10.1016/S1474-4422(15)70016-5

4. Selkoe DJ, Hardy J. The amyloid hypothesis of Alzheimer’s disease at 25 years. EMBO Mol Med. 2016;8(6):595–608. doi:10.15252/emmm.201606210

5. Hampel H, Hardy J, Blennow K, et al. The Amyloid-β Pathway in Alzheimer’s Disease. Mol Psychiatry. 2021;26(10):5481–5503. doi:10.1038/s41380-021-01249-0

6. Heneka MT, van der Flier WM, Jessen F, et al. Neuroinflammation in Alzheimer disease. Nat Rev Immunol. 2025;25(5):321–352. doi:10.1038/s41577-024-01104-7

7. Deczkowska A, Keren-Shaul H, Weiner A, Colonna M, Schwartz M, Amit I. Disease-Associated Microglia: A Universal Immune Sensor of Neurodegeneration. Cell. 2018;173(5):1073–1081. doi:10.1016/j.cell.2018.05.003

8. Keren-Shaul H, Spinrad A, Weiner A, et al. A Unique Microglia Type Associated with Restricting Development of Alzheimer’s Disease. Cell. 2017;169(7):1276–1290.e17. doi:10.1016/j.cell.2017.05.018

9. Yoo HJ, Kwon MS. Aged Microglia in Neurodegenerative Diseases: Microglia Lifespan and Culture Methods. Front Aging Neurosci. 2022;13:766267. doi:10.3389/fnagi.2021.766267

10. Götz J, Bodea LG, Goedert M. Rodent models for Alzheimer disease. Nat Rev Neurosci. 2018;19(10):583–598. doi:10.1038/s41583-018-0054-8

11. Sasaguri H, Hashimoto S, Watamura N, et al. Recent Advances in the Modeling of Alzheimer’s Disease. Front Neurosci. 2022;16. doi:10.3389/fnins.2022.807473

12. Saito T, Matsuba Y, Mihira N, et al. Single App knock-in mouse models of Alzheimer’s disease. Nat Neurosci. 2014;17(5):661–663. doi:10.1038/nn.3697

13. Stine WB, Dahlgren KN, Krafft GA, LaDu MJ. In vitro characterization of conditions for amyloid-beta peptide oligomerization and fibrillogenesis. J Biol Chem. 2003;278(13):11612–11622. doi:10.1074/jbc.M210207200

14. Kniewallner KM, Wenzel D, Humpel C. Thiazine Red+ platelet inclusions in Cerebral Blood Vessels are first signs in an Alzheimer’s Disease mouse model. Sci Rep. 2016;6(1):28447. doi:10.1038/srep28447

15. Mena R, Edwards P, P◆rez-Olvera O, Wischik CM. Monitoring pathological assembly of tau and ?-amyloid proteins in Alzheimer’s disease. Acta Neuropathol (Berl). 1995;89(1):50–56. doi:10.1007/BF00294259

16. Zhong MZ, Peng T, Duarte ML, Wang M, Cai D. Updates on mouse models of Alzheimer’s disease. Mol Neurodegener. 2024;19(1):23. doi:10.1186/s13024-024-00712-0

17. Mattsson-Carlgren N, Salvadó G, Ashton NJ, et al. Prediction of Longitudinal Cognitive Decline in Preclinical Alzheimer Disease Using Plasma Biomarkers. JAMA Neurol. 2023;80(4):360–369. doi:10.1001/jamaneurol.2022.5272

18. Neddens J, Temmel M, Flunkert S, et al. Phosphorylation of different tau sites during progression of Alzheimer’s disease. Acta Neuropathol Commun. 2018;6(1):52. doi:10.1186/s40478-018-0557-6

19. Masuda A, Kobayashi Y, Kogo N, Saito T, Saido TC, Itohara S. Cognitive deficits in single App knock-in mouse models. Neurobiol Learn Mem. 2016;135:73–82. doi:10.1016/j.nlm.2016.07.001

20. Portal B, Södergren M, Parés I Borrell T, et al. Early Astrocytic Dysfunction Is Associated with Mistuned Synapses as well as Anxiety and Depressive-Like Behavior in the *App ^NL^* ^-^ *^F^* Mouse Model of Alzheimer’s Disease. J Alzheimer’s Dis. 2024;100(3):1017–1037. doi:10.3233/JAD-231461

21. Shah D, Latif-Hernandez A, De Strooper B, et al. Spatial reversal learning defect coincides with hypersynchronous telencephalic BOLD functional connectivity in APPNL-F/NL-F knock-in mice. Sci Rep. 2018;8(1):6264. doi:10.1038/s41598-018-24657-9

22. Jankowsky JL, Zheng H. Practical considerations for choosing a mouse model of Alzheimer’s disease. Mol Neurodegener. 2017;12(1):89. doi:10.1186/s13024-017-0231-7

23. Sasaguri H, Nilsson P, Hashimoto S, et al. APP mouse models for Alzheimer’s disease preclinical studies. EMBO J. 2017;36(17):2473–2487. doi:10.15252/embj.201797397

24. Hemonnot-Girard AL, Valverde AJ, Hua J, et al. Analysis of CX3CR1 haplodeficiency in male and female APPswe/PSEN1dE9 mice along Alzheimer disease progression. Brain Behav Immun. 2021;91:404–417. doi:10.1016/j.bbi.2020.10.021

25. Dufour A, Heydari Olya A, Foulon S, et al. Neonatal inflammation impairs developmentally-associated microglia and promotes a highly reactive microglial subset. Brain Behav Immun. 2025;123:466–482. doi:10.1016/j.bbi.2024.09.019

26. Zappia L, Oshlack A. Clustering trees: a visualization for evaluating clusterings at multiple resolutions. GigaScience. 2018;7(7):giy083. doi:10.1093/gigascience/giy083

27. Aran D, Looney AP, Liu L, et al. Reference-based analysis of lung single-cell sequencing reveals a transitional profibrotic macrophage. Nat Immunol. 2019;20(2):163–172. doi:10.1038/s41590-018-0276-y

28. Silvin A, Uderhardt S, Piot C, et al. Dual ontogeny of disease-associated microglia and disease inflammatory macrophages in aging and neurodegeneration. Immunity. 2022;55(8):1448–1465.e6. doi:10.1016/j.immuni.2022.07.004

29. Neidert N, von Ehr A, Zöller T, Spittau B. Microglia-Specific Expression of Olfml3 Is Directly Regulated by Transforming Growth Factor β1-Induced Smad2 Signaling. Front Immunol. 2018;9. doi:10.3389/fimmu.2018.01728

30. Butovsky O, Jedrychowski MP, Moore CS, et al. Identification of a unique TGF-β–dependent molecular and functional signature in microglia. Nat Neurosci. 2014;17(1):131–143. doi:10.1038/nn.3599

31. Joshi JC, Joshi B, Zhang C, et al. RGS2 is an innate immune checkpoint for suppressing Gαq-mediated IFNγ generation and lung injury. iScience. 2025;28(2):111878. doi:10.1016/j.isci.2025.111878

32. Bibby JA, Agarwal D, Freiwald T, et al. Systematic single-cell pathway analysis to characterize early T cell activation. Cell Rep. 2022;41(8):111697. doi:10.1016/j.celrep.2022.111697

33. Luo OJ, Lei W, Zhu G, et al. Multidimensional single-cell analysis of human peripheral blood reveals characteristic features of the immune system landscape in aging and frailty. Nat Aging. 2022;2(4):348–364. doi:10.1038/s43587-022-00198-9

34. Barthélemy NR, Li Y, Joseph-Mathurin N, et al. A soluble phosphorylated tau signature links tau, amyloid and the evolution of stages of dominantly inherited Alzheimer’s disease. Nat Med. 2020;26(3):398–407. doi:10.1038/s41591-020-0781-z

35. Naia L, Shimozawa M, Bereczki E, et al. Mitochondrial hypermetabolism precedes impaired autophagy and synaptic disorganization in App knock-in Alzheimer mouse models. Mol Psychiatry. 2023;28(9):3966–3981. doi:10.1038/s41380-023-02289-4

36. Rahmani R, Rambarack N, Singh J, Constanti A, Ali AB. Age-Dependent Sex Differences in Perineuronal Nets in an APP Mouse Model of Alzheimer’s Disease Are Brain Region-Specific. Int J Mol Sci. 2023;24(19):14917. doi:10.3390/ijms241914917

37. Trojan E, Curzytek K, Cieślik P, et al. Prenatal stress aggravates age-dependent cognitive decline, insulin signaling dysfunction, and the pro-inflammatory response in the APPNL-F/NL-F mouse model of Alzheimer’s disease. Neurobiol Dis. 2023;184:106219. doi:10.1016/j.nbd.2023.106219

38. Andrews SJ, Renton AE, Fulton-Howard B, Podlesny-Drabiniok A, Marcora E, Goate AM. The complex genetic architecture of Alzheimer’s disease: novel insights and future directions. EBioMedicine. 2023;90:104511. doi:10.1016/j.ebiom.2023.104511

39. Ennerfelt H, Frost EL, Shapiro DA, et al. SYK coordinates neuroprotective microglial responses in neurodegenerative disease. Cell. 2022;185(22):4135–4152.e22. doi:10.1016/j.cell.2022.09.030

40. Schafer DP, Stillman JM. Microglia are SYK of Aβ and cell debris. Cell. 2022;185(22):4043–4045. doi:10.1016/j.cell.2022.09.043

41. Wang S, Sudan R, Peng V, et al. TREM2 drives microglia response to amyloid-β via SYK-dependent and -independent pathways. Cell. 2022;185(22):4153–4169.e19. doi:10.1016/j.cell.2022.09.033

42. Fu J, Ferreira D, Smedby Ö, Moreno R. Decomposing the effect of normal aging and Alzheimer’s disease in brain morphological changes via learned aging templates. Sci Rep. 2025;15(1):11813. doi:10.1038/s41598-025-96234-w

43. Rachmian N, Medina S, Cherqui U, et al. Identification of senescent, TREM2-expressing microglia in aging and Alzheimer’s disease model mouse brain. Nat Neurosci. 2024;27(6):1116–1124. doi:10.1038/s41593-024-01620-8

44. Jaganathan R, Iyaswamy A, Krishnamoorthi S, et al. Current aspects of targeting cellular senescence for the therapy of neurodegenerative diseases. Front Aging Neurosci. 2025;17:1627921. doi:10.3389/fnagi.2025.1627921

45. Findley CA, McFadden SA, Hill T, et al. Sexual dimorphism, altered hippocampal glutamatergic neurotransmission, and cognitive impairment in APP knock-in mice. J Alzheimers Dis JAD. 2024;102(2):491–505. doi:10.3233/JAD-240795

46. Izumi H, Shinoda Y, Saito T, et al. The Disease-modifying Drug Candidate, SAK3 Improves Cognitive Impairment and Inhibits Amyloid beta Deposition in App Knock-in Mice. Neuroscience. 2018;377:87–97. doi:10.1016/j.neuroscience.2018.02.031

47. Hebert LE, Weuve J, Scherr PA, Evans DA. Alzheimer disease in the United States (2010-2050) estimated using the 2010 census. Neurology. 2013;80(19):1778–1783. doi:10.1212/WNL.0b013e31828726f5

48. Viña J, Lloret A. Why women have more Alzheimer’s disease than men: gender and mitochondrial toxicity of amyloid-beta peptide. J Alzheimers Dis JAD. 2010;20 Suppl 2:S527–533. doi:10.3233/JAD-2010-100501

49. Guneykaya D, Ivanov A, Hernandez DP, et al. Transcriptional and Translational Differences of Microglia from Male and Female Brains. Cell Rep. 2018;24(10):2773–2783.e6. doi:10.1016/j.celrep.2018.08.001

50. Hao Y, Stuart T, Kowalski MH, et al. Dictionary learning for integrative, multimodal and scalable single-cell analysis. Nat Biotechnol. 2024;42(2):293–304. doi:10.1038/s41587-023-01767-y

51. Zhao Y, Fang ZY, Lin CX, Deng C, Xu YP, Li HD. RFCell: A Gene Selection Approach for scRNA-seq Clustering Based on Permutation and Random Forest. Front Genet. 2021;12. doi:10.3389/fgene.2021.665843

